# Stat2 loss disrupts damage signalling and is protective in acute pancreatitis

**DOI:** 10.1101/770750

**Authors:** Helen Heath, Gary Britton, Hiromi Kudo, George Renney, Malcolm Ward, Robert Hutchins, Graham R. Foster, Robert Goldin, William Alazawi

## Abstract

Severity of sterile inflammation, as seen in acute pancreatitis, is determined by damage-sensing receptors, signalling cascades and cytokine production. Stat2 is a type I interferon signalling mediator that also has interferon-independent roles in murine lipopolysaccharide-induced NF-κB-mediated sepsis. However its role in sterile inflammation is unknown. We hypothesised that Stat2 determines severity of non-infective inflammation in the pancreas.

Wild type (WT) and Stat2^−/−^ mice were injected intraperitoneally with cerulein or L-arginine. Specific cytokine-blocking antibodies were used in some experiments. Pancreata and blood were harvested 1h and 24h after the final dose of cerulein and up to 96h post L-arginine. Whole-tissue phosphoproteomic changes were assessed using label-free mass spectrometry. Tissue-specific Stat2 effects were studied in WT/*Stat2*^−/−^ bone-marrow chimera and using Cre-lox recombination to delete Stat2 in pancreatic and duodenal homeobox 1(*Pdx1*)-expressing cells.

*Stat2*^−/−^ mice were protected from cerulein- and L-arginine-induced pancreatitis. Protection was independent of type I interferon signalling. *Stat2*^−/−^ mice had lower cytokine levels including TNFα and IL-10 and reduced NF-kB nuclear localisation in pancreatic tissue compared to WT. Inhibition of TNFα improved (inhibition of IL-10 worsened) cerulein-induced pancreatitis in WT but not *Stat2*^−/−^ mice. Phosphoproteomics showed down-regulation of mitogen-activated protein kinase (MAPK) mediators but accumulation of Ser412-phosphorylated Tak1. Stat2 deletion in *Pdx1*-expressing acinar cells (*Stat2*^*flox*/*Pdx1-cre*^) reduced pancreatic TNFα expression, but not histological injury or serum amylase. WT/*Stat2*^−/−^ bone-marrow chimera were protected from pancreatitis irrespective of host or recipient genotype.

Stat2 loss results in disrupted signalling in pancreatitis, upstream of NF-κB in non-acinar and/or bone marrow derived cells.

## INTRODUCTION

The response to tissue damage involves signalling pathways such as Toll-like receptors, mitogen activated protein (MAP) kinases and transcription factors such as NF-κB [1]. Transcription factor activation initiates inflammatory and healing processes by producing pro- and anti-inflammatory cytokines that include TNFα, interleukins and interferons (IFN), which also serve to amplify the inflammatory response through their own signalling pathways. Stat2 is a mediator of type I IFN signalling, but it is increasingly evident that Stat2 has a key role in mediating inflammatory responses beyond this pathway [2, 3]. We have previously demonstrated an IFN-independent role for Stat2, but not other Stat proteins, in the response to bacteria- and virus-derived molecular patterns and in a mouse model of sepsis [4]. However, its role in non-infective inflammation, which is a characteristic of acute pancreatitis, has received scant attention to date.

Acute pancreatitis (AP) is a clinically significant example of damage-induced or ‘sterile’ inflammation. It is a potentially life-threatening, highly unpredictable and increasingly common inflammatory disease [5, 6] with no specific treatment. Production and activity of inflammatory cytokines are central to the pathogenesis of AP [7], but a number of challenges exist to studying acute pancreatitis in humans, especially in the early, initiating stages of disease. These include the inaccessibility of pancreatic tissue and a variable time interval between onset of symptoms and presentation to hospital. Mouse models can be used to address both of these challenges, but no single model recapitulates all the features of acute pancreatitis seen in man. Injection with supraphysiological concentrations of cerulein activates intracellular signalling and metabolic pathways that generate toxic mediators that cause a rapid mild pancreatitis. Injection with L-arginine causes a much more severe inflammation and can recapitulate later or secondary events in human pancreatitis.

To date, studies of type I IFN in the pathogenesis of pancreatitis have focused on known canonical functions of the type I IFN receptor (Ifnar) and signalling [8], but not Stat2. In the current study, we tested the hypothesis that Stat2-mediated signalling plays a role in AP as a model of non-infective tissue inflammation.

We have found that Stat2 loss is protective against pancreatic injury in two independent models of AP in mice. Stat2 loss was associated with reduced expression of inflammatory cytokine, and inhibiting cytokines can modulate the severity of pancreatitis in WT but not *Stat2*^−/−^ mice. We have used bone-marrow chimera and acinar cell-specific deletion of Stat2 to show that, despite the importance of acinar cells in cytokine production, non-acinar stromal and / or bone-marrow-derived cells determine the severity of tissue inflammation. This gives us novel insight into the role of Stat2 in mediating the severity of inflammation.

## MATERIALS AND METHODS

### Experimental models of AP

Generation of *Stat2*^−/−^ mice was previously described [9]. Mice were housed in a conventional facility in individually ventilated cages with free access to food and water. All animal studies were performed in accordance with the National Institutes of Health Guide for Care and Use of Laboratory Animals and approved the UK Home Office and Local Ethics Committee at Queen Mary, University of London. Patients were not involved in the design of this study.

Pancreatitis was induced in male and female mice age 6-9 weeks by hourly intraperitoneal injections of 50μg/kg or 100 μg/kg cerulein in PBS for 4h or 7h and samples were collected at 1h (hyperacute) and 24h (acute) after the last injection respectively. L-arginine-induced pancreatitis was induced by 2 intraperitoneal injections of 4g/kg L-arginine in PBS 1h apart.

For some experiments, mice were injected intraperitoneally with 100μg anti-TNFα (Ultra-LEAF™ Purified anti-mouse TNFα Antibody, clone MP6-XT22 Biolegend) or 200μg intraperitoneally (ip) anti-IL10 (LEAF™ purified anti-mouse IL10 Antibody, clone JES5-16E3 Biolegend) 1h before cerulein treatment. For IFNAR1 blocking experiments, mice were injected with 500μg ip anti-Mouse IFNAR1 Low Endotoxin Functional Formulation (clone MAR1-5A3 Universal Biologicals) 24hr before cerulein. For TNFα stimulation, mice received intraperitoneal injections of 2μg ip TNFα (Peprotech) 24h before cerulein injections. Control mice were injected with PBS alone. Blood samples were taken and serum was stored in aliquots at −80°C until use. Tissues were snap frozen for protein and RNA extraction or fixed immediately in 10% formalin and embedded in paraffin blocks for H&E and immunohistochemistry. Serum amylase levels were measured using a Roche Cobas analyser. Tissues were dried following incubation at 56°C for 72h. Tissue oedema was calculated by subtracting dry weight from initial wet weight.

### Serum Cytokine ELISA

Serum was analysed for TNFα, IFNγ, IL1β, IL6, IL10, IL17A using a multiplex (6-plex) ELISA (BD), and for IL-1B, IL-6 and IL-10 using the mouse magnetic luminex assay (R&D). Serum was used as 1:4 dilution in PBS for analysis.

### Quantitative Real-Time PCR

Tissue samples were sonicated on ice in Trizol (Thermo Fisher) and total RNA was isolated as recommended by the manufacturer. RNA concentrations were measured using NanoDrop 2000 (Thermo Scientific) and normalised across the experiment. cDNA was reverse transcribed using the Superscript IV protocol (Invitrogen) and qRT-PCR was performed using the rotor-gene SYBR green PCR kit (Qiagen) (see **Supplemental Table 1** for gene names and sequences used). Cycle threshold (ct) values were determined for each sample and the ct value difference (Δct) was used to calculate the factor of differential expression (2Δct) (Schmittgen and Livak, 2008). Correction for cDNA input deviations was performed using beta actin.

### Protein Immunoblots

Whole cell extracts were prepared by homogenising tissues in lysis buffer (Tris 50mM, NaCl 150mM, EDTA 2mM, 1% NP-40) containing Halt™ Protease and Phosphatase Inhibitor Cocktail (Thermo Fisher Scientific). Tissue samples were homogenised on ice, and incubated with vigorous shaking at 4°C for 30min. Cells were then vortexed and centrifuged for 10min at 13600rpm at 4°C. Cytoplasmic and nuclear extracts were prepared. Frozen tissues were homogenized in hypotonic buffer A (HEPES 10 mM, pH 7.5, KCl 10 mM, EGTA 0.1 mM, EDTA 0.1 mM, pH 8.0, 1 mM DTT) containing protease/phosphatase inhibitor cocktail. Cells were kept on ice for 10 minutes, 0.3% NP-40 added, vortexed briefly and spun down for 30s at 4°C. Supernatant (cytoplasmic fraction) was removed and stored on ice for protein quantification. The pellet was washed again with buffer A, and the hypertonic, high-salt buffer C was added (HEPES 20 mM, pH 7.5, NaCl 0.4 M, EDTA 1 mM, EGTA 1 mM, glycerol 20%) together with the protease/phosphatase inhibitor cocktail. After centrifugation (10 min at 13600rpm, 4°C), the supernatant was collected and referred to as the nuclear fraction.

Supernatants from whole cell lysis and fractionated extracts were collected and protein measurements were done using the Pierce BCA assay as recommended by the manufacturer (Thermo Fisher). Aliquots of equal concentration were mixed in 4X sample buffer consisting of 277.8 mM Tris-HCl, pH 6.8, 44.4% (v/v) glycerol, 4.4% LDS, 0.02% bromophenol blue (Bio-Rad) with additional 50mM DTT, boiled at 95°C for 5min and stored at −80°C. See **Supplemental Table 1** for antibodies used.

### Histology/immunocytochemistry

Sections of formalin fixed paraffin embedded tissues were stained with haematoxylin and eosin and analysed on a histology microscope. A pathologist, blinded to the treatment, group scored, digitised, images of the whole specimen for oedema (0, absent or rare; 1, oedema in the interlobular space; 2, oedema in the intralobular space; and 3, the isolated-island shape of pancreatic acinus), inflammation (0, absent; 1, mild; 2, moderate; and 3, severe) and parenchymal necrosis (0, absent; 1, focal (< 5%); 2, and/or sublobular (< 20%); 3, and/or lobular (> 20%)). parenchymal haemorrhage was not seen. Images were taken on a Hammamatsu scanner and displayed at 20x magnification.

### Immunofluorescence Microscopy

Pancreas cell suspensions were prepared as described [10]. Isolated acinar cells were plated onto chamber slides (NUNC) pre-coated with 50 μg/ml type 1 collagen (Thermo Fisher) in 0.02 M acetic acid, 0.2 μm-filtered and incubated overnight at 37°C 5% CO_2_. Cells were treated with 10ng/ml TNFα (Peprotech) or 1nM CCK (Cambridge Biosciences) for indicated time points. Cells were fixed in cold PFA (4% PFA in PBS) for 20min room temperature. Cells were permeabilised in 0.3% Triton X100 in PBS for 10min and blocked in 5% BSA/PBS for 30min at RT. Cells were incubated with anti-p65 (D14E12) XP^®^ Rabbit mAb CST) primary antibody (1:200) in 2% BSA/PBS/0.05% Tween-20 at 4°C overnight. See **Supplemental Table 1** for antibodies used. Cells were washed 3X with wash buffer (PBS/0.05% Tween-20) and stained with goat anti-rabbit 488 conjugated antibody 1:500 (Thermo Fisher) in in 2% BSA/PBS/0.05% Tween-20 for 1h at RT. Slides were washed in wash buffer 2X and PBS 1X and dehydrated in 70-100% ethanol series prior to mounting in VECTASHIELD Antifade Mounting Medium with DAPI (Vector Laboratories). Confocal images were acquired on Zeiss LSM 710 upright scanning confocal microscope using 63X oil objective, numerical aperture 1.4. Z-stacks were acquired at 1μm intervals 20μm above and below the midline of the cell cluster. Images were analysed using ImageJ and IMARIS (Bitplane) software. Localisation of indirectly labelled p65 in the nuclei was quantified by preparing a 3-D threshold image in DAPI and measuring the mean intensity values for the green channel in each nuclei.

### Statistics

All serum amylase, ELISA, IMARIS and real-time PCR data were analysed using Prism software. Data are presented as mean with error bars reflecting standard error of the mean (SEM). Mass spectrometry quantitative normalised total spectra for phosphorylation modifications were analysed using Scaffold Viewer 4.7.2 (Proteome Software, Portland, Oregon). Pathway analysis was performed using PANTHER gene enrichment software (http://pantherdb.org).

## RESULTS

### *Stat2*^−/−^ mice are protected from acute pancreatitis

To assess the effect of Stat2 on tissue inflammation, AP was induced in WT and *Stat2*^−/−^ mice by giving hourly intraperitoneal (i.p) injections of cerulein for 4h or 7h. Tissue and blood samples were harvested 1h (which we defined as hyperacute) and 24h (which we defined as acute) after the final injection respectively (**Supplemental Fig. 1a**). Stat2^−/−^ mice had significantly lower serum levels of amylase compared to WT following hyperacute injury (**Fig. 1a**, 3808iU/ml vs 8001iU/ml, p<0.0001). At this time point, pancreatic tissue damage was mild, but there was significantly less pathology (in particular oedema) in *Stat2*^−/−^ mice compared to WT (**Fig. 1b** **and Supplemental Fig. 1b**). In the acute model (7 hourly injections of cerulein and harvesting after 24h), injury was more evident histologically and *Stat2*^−/−^ mice had less pancreatic necrosis and interlobular oedema compared to WT (**Fig. 1b**), although amylase levels had returned to baseline in both genotypes at this time point (not shown).

**Figure 1.**
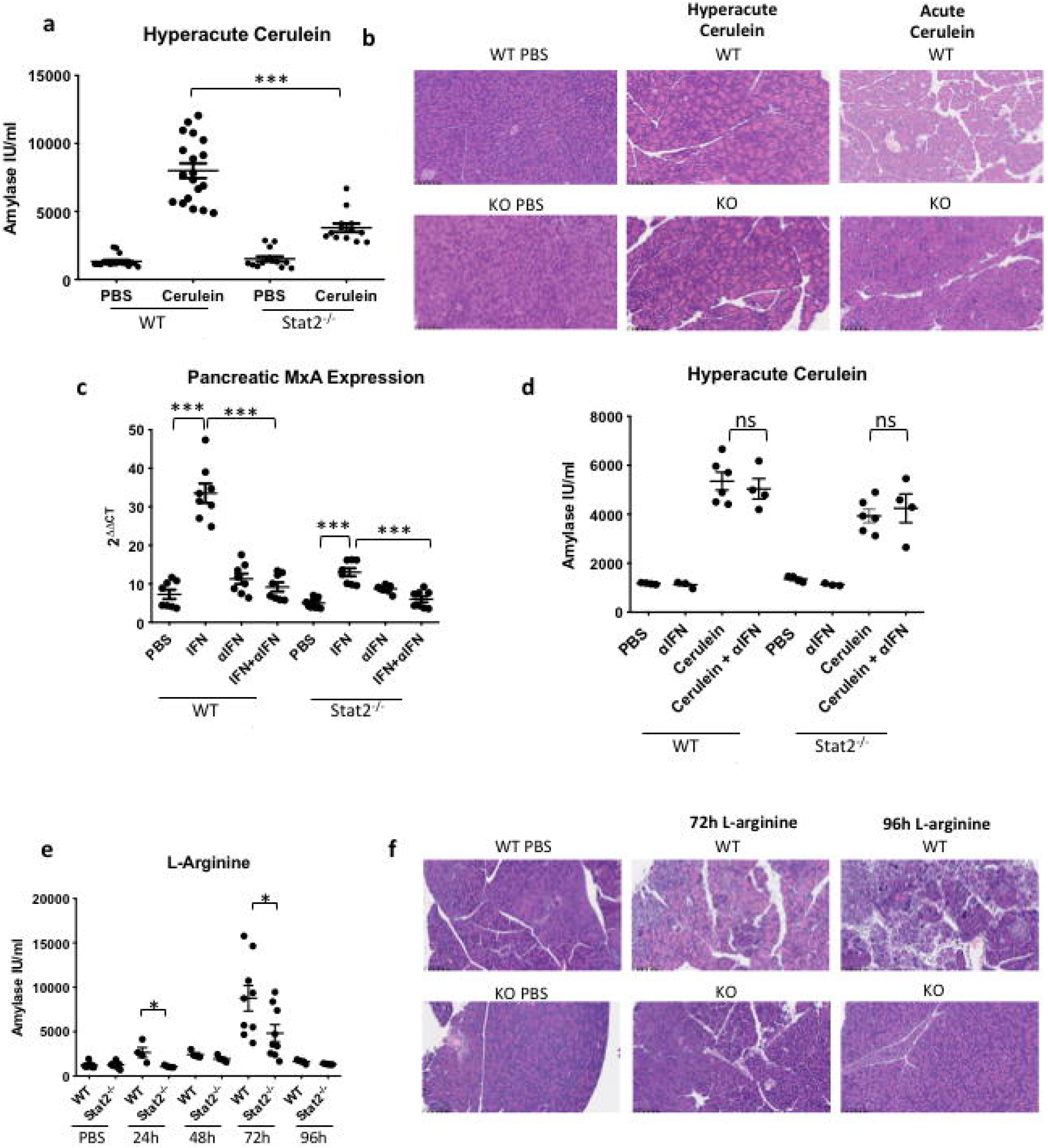
Stat2 loss reduces the severity of experimental pancreatitis. (**a**) Serum concentration of amylase and (**b**) representative haematoxylin and eosin-stained histology following injection with ceruelin at the time points indicated. Effect of Ifnar1 blocking antibody on (**c**) IFNα-induced Mx1 gene expression and (**d**) cerulein-induced serum amylase expression. (B) Serum concentration of amylase and (**f**) histology following injection with L-arginine at the time points indicated. WT – wild type, KO - Stat2^−/−^, PBS – phosphate buffered saline, Cer – cerulein ***p<0.001, *p<0.05, ns – not significant.

We did not find any differences in the expression of the receptors that are activated by cerulein (cholecystokinin (CCK)-AR or CCK-BR) between WT and *Stat2*^−/−^ pancreatic extracts that could account for the difference in severity of injury observed (**Supplemental Fig 1c**). Total Stat2 levels in whole cell extracts from whole WT pancreata were unaffected by cerulein treatment (**Supplemental Fig 1d**). We excluded the possibility that loss of Stat2 protects from hyperacute cerulein-induced pancreatitis through a type I IFN-mediated signalling loop by pre-treating animals with 500μg/i.p Ifnar-1 blocking antibody. Although the antibody effectively inhibited IFNα-mediated Mx1 production in splenocytes (**Fig 1e**), it had no significant impact on amylase levels in *Stat2*^−/−^ mice and did not protect WT mice from cerulein-induced pancreatitis (**Fig 1f**).

Stat2 loss was protective in a second, more severe model of pancreatitis induced by L-arginine injection. *Stat2*^−/−^ mice had lower levels of serum amylase compared to WT mice at 24h and 72h following L-arginine treatment (8768iU/ml vs 4829iU/ml, p<0.05, **Fig 1e**). L-arginine caused marked pancreatic necrosis, oedema, tissue destruction and inflammatory cell infiltration over 96h in WT mice, that was much less evident in *Stat2*^−/−^ mice at all time points (**Fig. 1f**, **Supplemental Fig. 1e-f**).

### Stat2 mediates acute pancreatitis upstream of cytokine production and activity

We did not observe clinical signs of systemic inflammation nor significant increases in serum concentrations of cytokines in hyperacute or acute cerulein-treated models (**Supplemental Fig. 2**). However, elevated serum IL-6, IL-1β and IL-10 were detected at later time points in WT L-arginine-treated mice (72h), and these levels were significantly lower in Stat2^−/−^ mice (**Fig 2a**).

**Figure 2.**
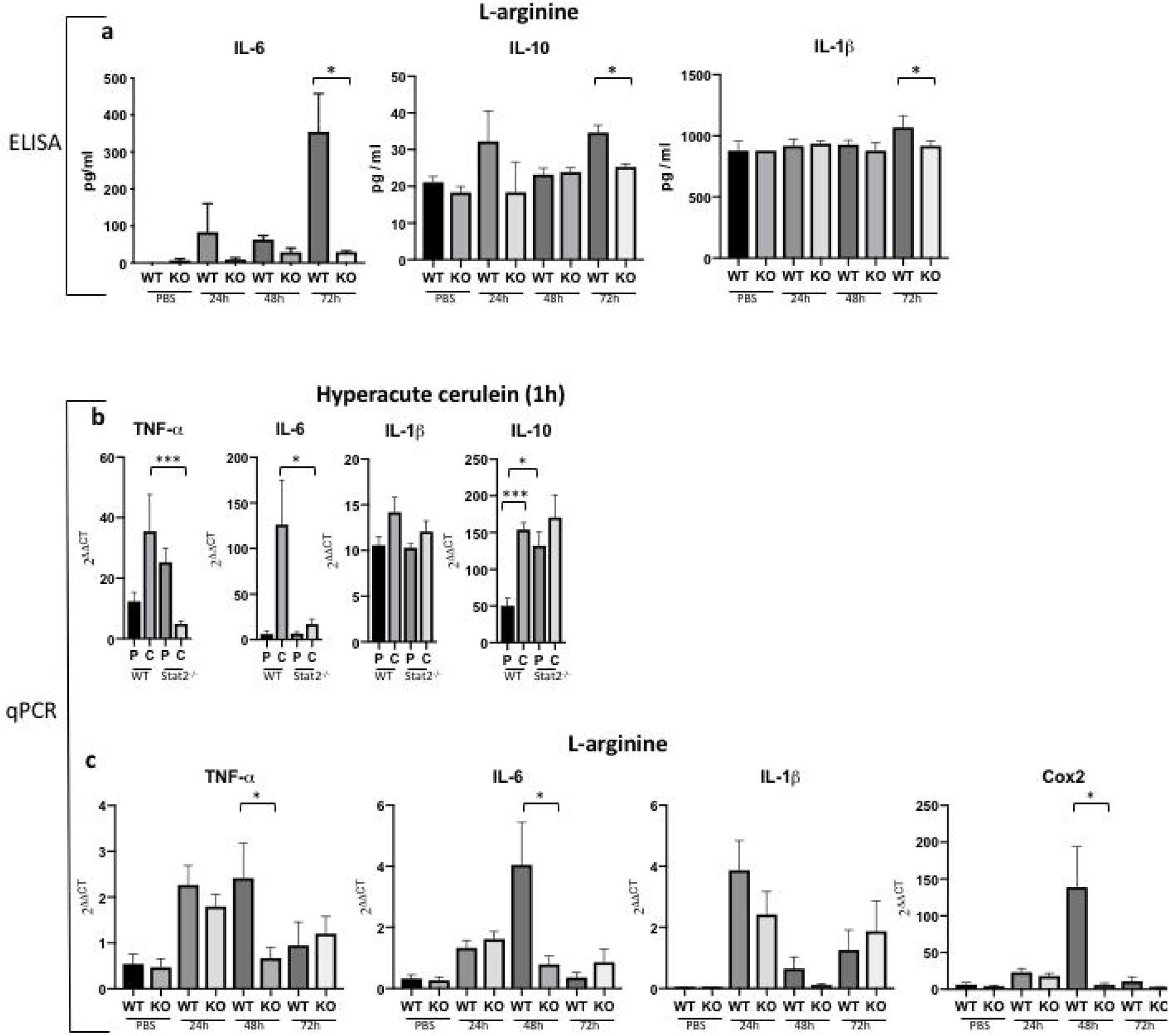
The effect of Stat2 loss on cytokine production. (**a**) Serum cytokine concentrations in wild type (WT) and Stat2^−/−^ (KO) mice following treatment with PBS or L-arginine. RNA expression of genes shown in wild type (WT) and Stat2^−/−^ (KO) whole pancreata following treatment with PBS (P) or cerulein (C) in (**b**) hyperacute cerulein- or (**c**) L-arginine-induced pancreatitis. *p<0.05.

Acute pancreatitis led to increased expression of inflammatory genes in pancreatic tissue. Consistent with protection from injury, expression of inflammatory genes, TNFα and IL-6, was lower in *Stat2*^−/−^ pancreas compared to WT (**Fig 2b**) in cerulein- and L-arginine-induced pancreatitis (**Fig 2c**). In contrast, the levels of the anti-inflammatory gene IL-10 were higher in PBS-treated control *Stat2*^−/−^ mice compared to WT and while treatment with cerulein increased the expression of IL-10 in WT mice, there was no further increase from baseline in *Stat2*^−/−^ mice. Therefore, we tested the hypothesis that inhibition of TNFα in mice would protect against acute pancreatitis, while inhibition of IL-10 would worsen tissue injury in WT mice. In order to study early inflammatory and not secondary events, we focused on the hyperacute cerulein model. Pre-administration of 100μg i.p TNFα blocking antibody resulted in significant reduction in serum amylase (6483iU/ml WT v 4442iU/ml WT+anti-TNFα, p<0.05) and histological injury in WT but not *Stat2*^−/−^ mice (**Fig. 3a-c**). Pre-treatment with recombinant TNFα (2μg i.p.) did not significantly increase serum amylase in cerulein-treated mice of either genotype (**Supplemental Fig. 3**). Pre-treatment with 200μg i.p IL-10-blocking antibody resulted in increased serum amylase (1997iU/ml WT v 2873iU/ml WT+anti-IL-10, p<0.05) and histological injury in WT mice, but did not have an effect in *Stat2*^−/−^ mice (**Fig. 3d-f**), suggesting that Stat2 acts upstream of production of cytokines such as TNFα or IL-10.

**Figure 3.**
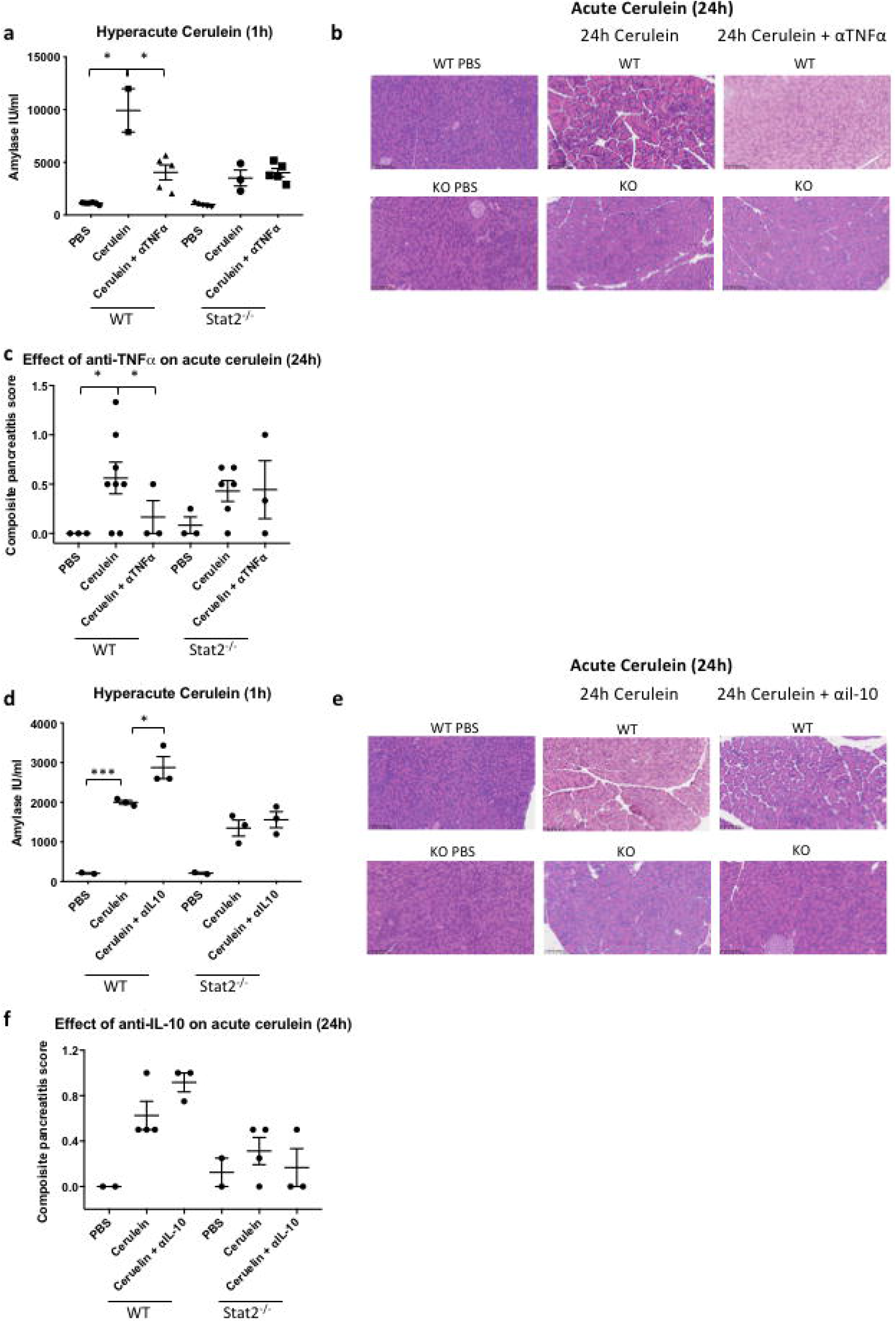
Effects of blocking TNFα and IL-10 on cerulein-induced pancreatitis. Wild type (WT) and Stat2^−/−^ (KO) mice were given a single intraperitoneal injection of (**a-c**) 100μg TNFα blocking antibody or (**d-f**) 200μg IL-10-blocking antibody prior to PBS or cerulean treatment according to the hyperacute protocol. (**a,d**) Serum amylase, (**b,e**) representative haematoxylin and eosin-stained histology and (**c,f**) tissue injury score are shown for each pre-treatment. *p<0.05

The NF-κB family of transcription factors mediates cytokine release following tissue injury. To model the activation of NF-κB in vitro, we stimulated primary acinar cells from WT or *Stat2*^−/−^ mice with CCK-8, a synthetic octapeptide analogue of cerulein [11,12]. We found a rapid and statistically significant shift from cytoplasmic to nuclear localisation of NF-κBp65 at 5 and 15min in WT cells, whereas in *Stat2*^−/−^ cells, there was little evidence of the oscillation of NF-κBp65 (**Fig. 4a-b**). Thus in pancreas tissue, Stat2 influences the cellular localisation of NF-κBp65 and hence cytokine production.

**Figure 4.**
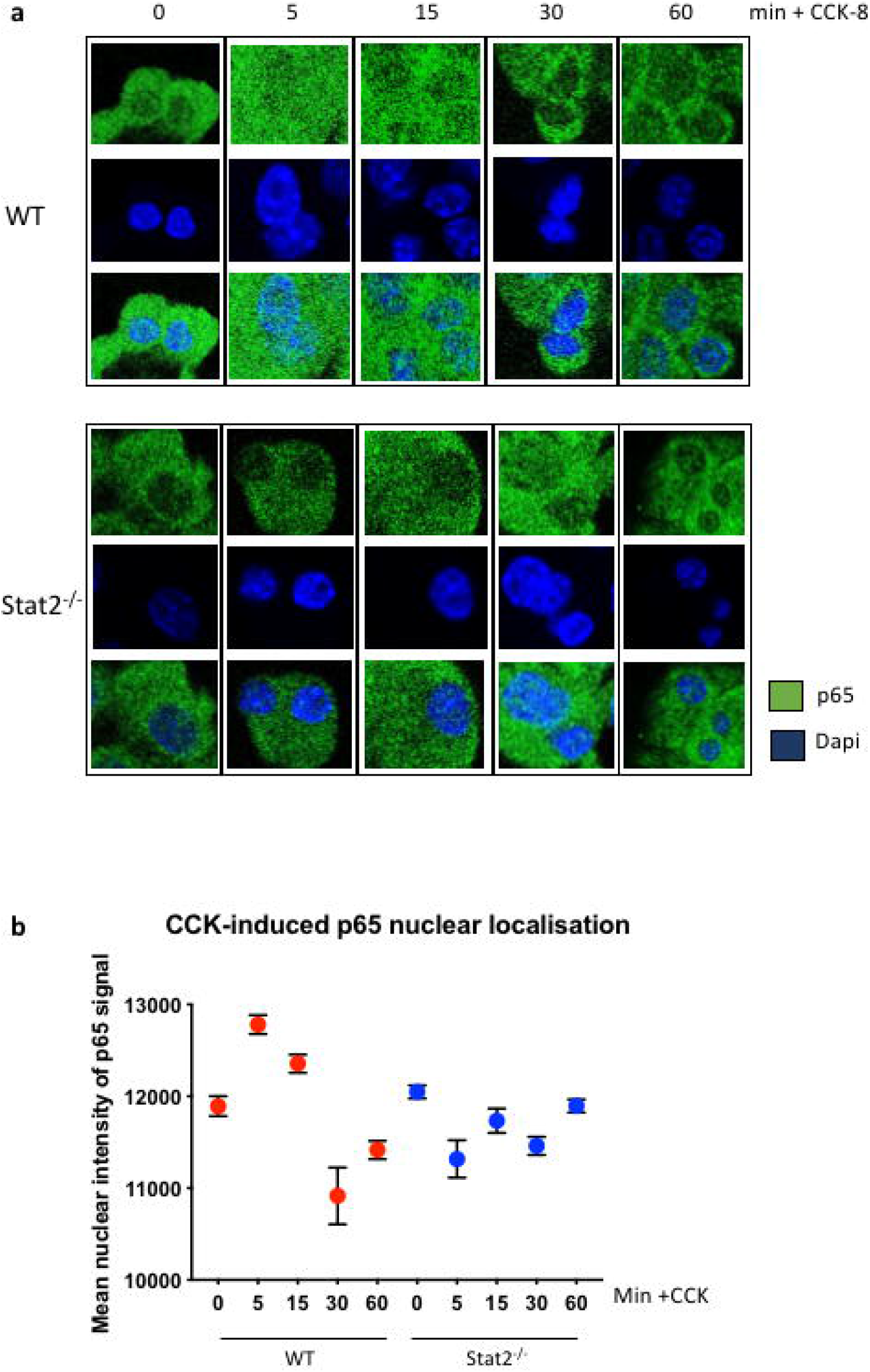
Loss of Stat2 disrupts NF-κB nuclear localisation. (**a**) Confocal microscopy showing NF-κBp65 staining (green) and nuclear DAPI (blue) in primary pancreatic cells at the timepoints shown following stimulation with CCK-8. (**d**) IMARIS software quantification of nuclear localization of NF-κBp65 signal in confocal images. WT – wild type, KO - Stat2^−/−^

### Loss of Stat2 impairs signalling in damage pathways

To determine whether Stat2 loss had a more global impact on intracellular signalling that leads to cytokine production, we used label-free tandem mass spectrometry to study Stat2- and cerulein-dependent phosphoprotein abundance within the pancreas. Total protein was extracted from WT and *Stat2*^−/−^ pancreata following treatment with either PBS or cerulein (hyperacute model, n=4 in each group) and processed for whole phosphoproteomic analysis as previously described [13]. We detected 611 proteins with peptide sequence concordance threshold 95% and protein threshold 99.9% (used to confirm the presence and identity of a protein). Based on the expression level of these proteins, 15 out of 16 samples clustered according to the experimental condition in an unsupervised hierarchical clustering algorithm (**Fig. 5a** **and Supplemental Fig. 4**). Overall the total number of phosphoproteins identified was comparable in each of the experimental conditions (**Fig. 5b**).

**Figure 5.**
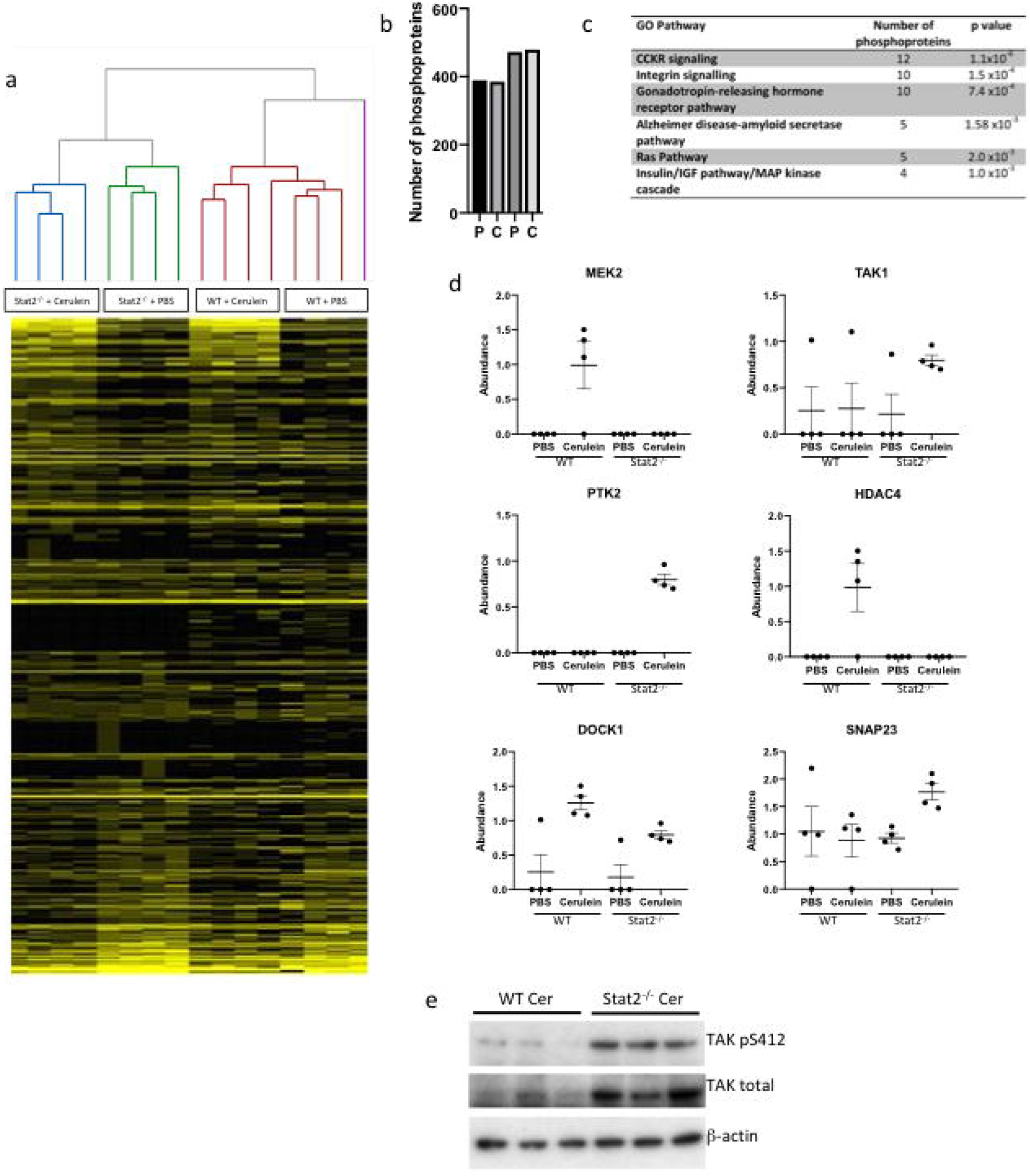
Global phosphoproteomic changes associated with Stat2 loss. 22,663 tandem mass spectra were identified, corresponding to 2881 unique peptide ions, of which 1650 were phosphopeptides containing 1771 phosphopeptide sites. (**a**) unsupervised hierarchical clustering of 611 phosphoproteins identified using label-free mass spectrometry. Each lane represents an individual mouse and each row is one phosphoprotein. Yellow indicates high abundance and black lowest abundance. (**b**) Total number of phosphoproteins identified in each experimental condition. WT – wild type, KO - Stat2^−/−^, PBS – phosphate buffered saline. (**c**) Signalling pathways identified by GO Panther pathway analysis to be statistically significantly enriched in 278 phosphoproteins that differed between WT and Stat2^−/−^ pancreata either at baseline or after treatment with cerulein. (**d**) Abundance of key signalling phosphoproteins as determined by mass spectrometry. (**e**) Western blot for phosphorylated and total Tak1 in whole cell extracts taken from wild type (WT) and Stat2^−/−^ whole pancreata following treatment with PBS or cerulein in the hyperacute model.

In order to maximise signalling pathway discovery, we used pre-determined thresholds for differences of interest at abundance ratios of <0.75 and >1.33. We identified 348 unique phosphoproteins that were differently expressed in our experimental conditions (**Supplemental Table 2**). The expression of 257 phosphoproteins was altered in WT pancreas as a result of cerulein treatment. In total 175 phosphoproteins were altered in cerulein-treated *Stat2*^−/−^ vs WT pancreas, of which 128 were detected in the inflamed but not control samples (66 upregulated and 62 downregulated in *Stat2*^−/−^). We conducted a GO Panther analysis of all 175 phosphoproteins that were differentially expressed between WT and *Stat2*^−/−^ in unstimulated and in stimulated tissue. This highlighted pathways involved in signalling and the response to injury, including the CCK receptor signalling pathway (n=12 phosphoproteins, p=1.1×10^−6^, **Fig. 5c**). Five out of 6 significantly identified pathways contain at least one mitogen-activated protein (MAP) kinases, including MAP2K2 (MEK2), the principal activator of extracellular signal related kinases (ERK1 and ERK2, [14]). The activating phosphorylation of MEK2 at serine S226 was more abundant in inflamed WT pancreata compared to Stat2^−/−^ in the phosphoproteome analysis by mass spectrometry (**Fig. 5d**). Using Western blotting, we found that the expression ERK1/2 pT202/Y204, a downstream target of MEK2, was also reduced in *Stat2*^−/−^ nuclear pancreatic extracts compared to WT (**Supplemental Fig. 5a**) [15].

While we observed no difference in two upstream activators of MEK2 (MKK3 pSer189 or MKK6 pSer207) between WT and *Stat2*^−/−^ pancreata (**Supplemental Fig. 5b**), we did find increased levels of Map3K7 (TGFβ-activated protein kinase 1: Tak1), a MAPKKK that lies upstream of MEK2 and NF-kBp65. The phosphorylated Ser412 modification of Tak1 was significantly more abundant in *Stat2*^−/−^ cerulein-treated samples compared to other conditions by mass spectrometry (**Fig. 5d**) and in independent experiments by western blot (**Fig 5e**). This suggests that Ser412-phosphorylation of Tak1 is a potential point of pathway interruption in the absence of Stat2.

### Cell-specific effects of Stat2 loss

To determine the cell type in which Stat2 plays a role, we generated bone marrow chimeric mice and found that loss of Stat2 in either donor or recipient cellular compartments resulted in lower serum amylase (**Fig. 6a**) and less histological injury (**Fig. 6b**). To further refine this, we generated mice with a conditional deletion of Stat2 (*Stat2*^*flox*/*flox*^) induced by Pdx1-cre to specifically delete Stat2 in cells expressing *Pdx1* (*Stat2*^*Δacinar*^) [16]. Following hyperacute cerulein-induced pancreatitis, the expression of inflammatory TNFα was significantly lower in *Stat2*^*Δacinar*^ mice compared to WT (**Fig. 6b**), in keeping with the impaired signalling above. However this had no significant effect on serum amylase concentration (**Fig. 6c**) nor pancreatic histology (**Fig. 6d**). Given that cytokine blockade did modulate pancreatic inflammation, these results strongly suggest that Stat2- and cytokine-dependent tissue injury in acute pancreatitis is mediated by a combination of bone-marrow-derived cells and/or non-acinar stromal cells.

**Figure 6.**
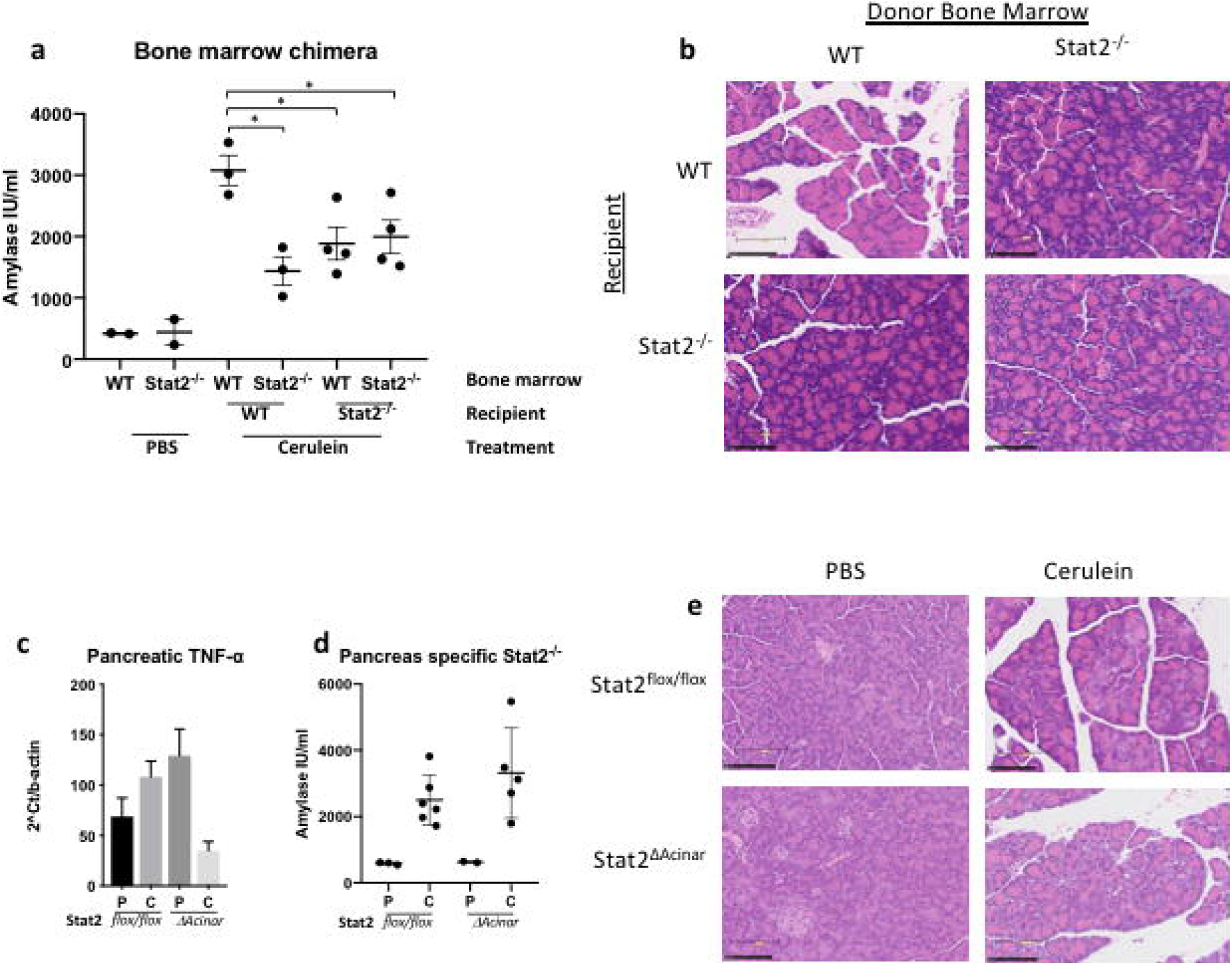
Effects of lineage Stat2 deletion in specific compartments on cerulein-induced pancreatitis. (**a**) Serum concentration of amylase and (**b**) representative haematoxylin and eosin-stained histology in bone-marrow chimeric mice following hyperacute cerulein-induced pancreatitis. (**c**) RNA expression of TNFα in whole pancreatic extracts, (**d**) serum concentration of amylase and (**b**) representative haematoxylin and eosin-stained histology in *Stat2*^*flox*/*Pdx1-cre*^ mice following hyperacute cerulein-induced pancreatitis.

## DISCUSSION

Our data show that Stat2 has a pathogenic, pro-inflammatory role in the response to sterile injury in the pancreas. In two models, assessed at different time points, Stat2^−/−^ mice were protected from cerulein- and L-arginine-induced acute pancreatitis. Using a phosphoproteomic approach, we identified signalling mediators from the MAPK cascade downstream of, and including, Tak1, that were differentially activated between WT and *Stat2*^−/−^. However, acinar cell-specific deletion of Stat2 alone is insufficient for histological and biochemical protection from acute pancreatitis, suggesting that Stat2 plays a role in non-acinar stromal and/or bone-marrow-derived cells.

Cerulein is believed to cause pancreatitis by supra-physiological activation of CCK1 and CCK2 receptors, which signal through MAPK intermediates [17]. We have shown that in vivo cytokine gene expression in response to cerulein is Stat2-dependent, as is NF-κB localisation to the nucleus in primary acinar cells. L-arginine does not act through CCK receptors, yet Stat2 loss is also protective in this model of pancreatitis. The mechanism by which L-arginine causes pancreatitis is not fully understood but cellular metabolic disturbances liberate damage-associated molecular patterns which can trigger MAPK-containing signalling pathways [18, 19]. Thus Stat2 seems to play a wide role in MAPK-mediated signal transduction in response to injury that is agnostic of the initial triggering stimulus.

The main question that arises from our work is where in these signalling pathways Stat2 is acting. The data show an effect on MAPK-mediated signalling, ERK activation and NF-κB signalling. One interpretation of our results is that Stat2 sits at a common point that is proximal in these pathways. A build-up of pS413Tak1 in inflamed pancreatic tissue in *Stat2*^−/−^ mice might suggest that Stat2 is downstream of Tak1 but we have not excluded a compensatory increase in kinases or phosphatases in the absence of Stat2 that affects upstream signalling. Similarly we have not demonstrated a direct interaction between Stat2 and Tak1 or other mediators of these pathways to see if there is physical association.

We could not recapitulate the protection seen in *Stat2*^−/−^ mice by blocking type I IFN signalling. Indeed, at the same early time points assessed in the current study, no difference in pancreatic damage was observed between WT and *Ifnar1*^−/−^ mice [20]. This suggests that IFN-mediated signals are not required for early pancreatitis and earlier work has shown that cellular responses to epidermal growth factor [21] and LPS [4] are mediated by Stat2 even in the absence of the IFN phosphorylation site at Tyr689. Previous studies into the role for type I IFN signalling in acute pancreatitis have used prolonged injection schedules and focussed on much later time points after the final injection (3-7 days). In these models, deletion of Ifnar1 partially improved tissue injury without an effect on cytokine mRNA levels [8, 20]. Conversely, we find that pancreas-specific deletion of Stat2 reduces TNFα levels but has no effect on tissue injury. Taken together, these data emphasise the IFN-independent role for Stat2 in AP and challenge the dogma that cytokine production is an obligate determinant of tissue injury. Nevertheless, we show that specific cytokine modulation does alter the severity of tissue inflammation and the main effect of Stat2 is to mediate signalling that leads to NF-κB activation.

A consensus has yet to be reached on whether NF- κB activation is protective or deleterious in acute pancreatitis [22, 23]. This is largely because NF- κB activation is the result of the timely integration of multiple pathways and NF- κB itself can activate pro- and anti-inflammatory downstream effectors [24]. Pancreas-specific mutants of NF-κBp65 that are unable to dimerise or translocate to the nucleus resulted in more severe cerulein-induced injury [25] and deletion of IκBa, which results in reduced activation of NF-κBp65 and reduced cerulein-induced acute pancreatitis [23]. Conversely, conditional overexpression of NF-κB or constitutively active IKK2 increased activity of NF-κB in acinar cells, and led to more severe pancreatitis compared to control mice [22]. Stat2 deficiency in our models is constitutive rather than inducible, but we do show that the absence of Stat2 is associated with disrupted nuclear entry and exit of NF-κBp65. This finding is complementary to our previous working showing a role for Stat2 in response to the pathogen-associated molecular pattern lipopolysaccharide; a model of infective inflammation [4]. We have not over-expressed Stat2, so do not know if this results in NF-κB over-activation nor the effects that this would have on severity. Our results suggest that at early time points, NF-κB related signalling and gene expression do contribute to disease severity hence early cytokine inhibition alters the disease course. The protective effect of TNFα inhibition is of clinical as well as pathogenic interest and a clinical trial of infliximab versus placebo in patients with acute severe pancreatitis is currently underway. Our data certainly support the hypothesis that early inhibition of TNFα can be beneficial in pancreatitis, but further work is needed to determine the characteristics of patients who will benefit and the timing of treatment to maximise efficacy and minimise potential side effects such as sepsis. We have shown that IFN-independent, Stat2-dependent signalling leads to the timely activation of NF-κB in models of sterile pancreatic inflammation. Strategies that target Stat2 or its associated signalling mediators may have a therapeutic role in acute sterile inflammatory disease.

## ACKNOWLEDGEMENTS

We thank Professor Christian Schindler for sharing the Stat2^−/−^ mice with our laboratory. We are grateful to Dr Pedro Cutillas who ran proteomic mass spectrometry assays, Polychronis Kemos for statistical assistance and Jan Soetaert and Belen Martin-Martin from the Blizard Advanced Light Microscopy facility.

## Author Contributions

HH, GB, HK, GR, MW, RG conducted experiments and/or analysed data

WA and GRF developed the study concept

WA and HH designed experiments

RH and GRF interpreted data and critically reviewed the manuscript

HH and WA wrote the paper and all authors reviewed and approved the paper

WA supervised the project

The authors declare that they have no conflict of interest

---

**Supplemental Figure 1.** (**a**) Mice were injected according to the schedules shown (solid arrows) and blood and tissues harvested (dotted arrows). (**b**) Oedema in pancreatic tissue demonstrated by net wet-dry weight. (**c&d**) In hyperacute cerulein experiments, Western blots showing (c) expression of cholecystokinin (CCK) receptors A and B and (**d**) of Stat2 in pancreata from WT and *Stat2*^−/−^ mice. (**e**) Composite issue injury score and (**f**) representative haematoxylin and eosin-stained histology in L-arginine-induced pancreatitis at time points indicated. P – treated with PBS, C – treated with cerulein.

**Supplemental Figure 2.** Serum cytokine concentrations in wild type (WT) and Stat2^−/−^ mice following treatment with PBS (P) or cerulein (C) at time points indicated.

**Supplemental Figure 3.** Effect of pre-treatment with TNFα on serum concentration of amylase hyperacute cerulein-induced pancreatitis.

**Supplemental Figure 4.** Principal component analysis of phosphoproteome identified by label-free mass spectrometry in whole pancreata from wild-type (WT) and *Stat2*^−/−^ (KO) mice following treatment with PBS or hyperacute cerulein-induced pancreatitis.

**Supplemental Figure 5.** Western blots showing abundance of proteins and phosphoproteins shown in (**a**) nuclear and cytoplasmic or (**b**) whole cell extracts from whole pancreata following hyperacute cerulein-induced pancreatitis.

