## Supplemental Table 1 for "Stat2 loss disrupts damage signalling and is protective in acute pancreatitis"

### Supplemental Table 1. Methods

#### A. Antibody list

| Antibody | Company and cat # | Use | dilution |
|---|---|---|---|
| p44/42 MAPK (Erk1/2) Antibody | CST 9102 | WB | 1:1000 |
| Phospho-p44/42 MAPK (Erk1/2) (Thr202/Tyr204) | CST 9101 | WB | 1:1000 |
| α-Tubulin (11H10) Rabbit mAb | CST 2125 | WB | 1:1000 |
| Anti-beta Actin | Abcam ab8224 | WB | 1:2500 |
| p38 MAPK (D13E1) XP® Rabbit mAb | CST 8690 | WB | 1:1000 |
| Phospho-p38 MAPK (Thr180/Tyr182) (D3F9) XP® Rabbit mAb | CST 4511 | WB | 1:1000 |
| Fibrillarin (C13C3) Rabbit mAb | CST 2639 | WB | 1:1000 |
| NF-κB p65 (D14E12) XP® Rabbit mAb | CST 8242 | WB, IF | 1:1000, 1:500 |
| IKKβ (D30C6) Rabbit mAb | CST 8943 | WB | 1:1000 |
| IKKα (3G12) Mouse mAb | CST 11930 | WB | 1:1000 |
| IκBα (L35A5) Mouse mAb | CST 4814 | WB | 1:1000 |
| NF-κB1 p105/p50 (D4P4D) Rabbit mAb | CST 13586 | WB | 1:1000 |
| Stat2 polyclonal antibody | Merck Millipore 07-140 | WB | 1:2000 |
| Stat2 (D9J7L) Rabbit mAb | CST 72604 | WB, IF | 1:1000, 1:500 |
| Anti-CCKAR/CCK1R antibody | Invivogen SourceBioscience LS-C177096 | WB | 1:500 |
| Anti CCKBR (E3) antibody | Santa Cruz sc-166690 | WB | 1:1000 |
| IL1B (D3H1Z) Rabbit mAb | CST 12507 | WB | 1:1000 |
| Phospho-TAK1 (Ser412) Antibody | CST 9339 | WB | 1:1000 |
| Phospho-MEK1/2 (Ser218/222/226) | Abcam ab78132 | WB | 1:500 |
| MKK3 (D4C3) Antibody | CST 8535 | WB | 1:1000 |
| MKK6 (D31D1) Antibody | CST 8550 | WB | 1:1000 |
| Phospho-MKK3/6 (D8E9) (Ser189/207) | CST 12280 | WB | 1:1000 |
| Phospho-c-Jun (Ser73) | CST 9164 | WB | 1:1000 |
| F(ab')2-Goat anti-Mouse IgG (H+L) Cross-Adsorbed Secondary Antibody, Alexa Fluor 488 | ThermoFisher A11017 | IF | 1:2000 |
| Peroxidase-AffiniPure F(ab')2 Fragment Donkey Anti-Rabbit IgG (H+L) | Jackson Immunoresearch 711-036-152-JIR | WB | 1:30000 |
| Anti-mouse IgG, HRP linked Antibody | GE NA931 | WB | 1:3000 |
| Anti-mouse IgG, HRP-linked Antibody | CST 7076 | WB | 1:1000 |

**B. Oligonulceotide sequencess**

| Name | Sequence 5'-3' | Use |
|---|---|---|
| mmTNFaF | CCAGTGTGGGAAGCTGTCTT | qRTPCR |
| mmTNFaR | AAGCAAAAGAGGAGGCAACA | qRTPCR |
| mmIL6F | GTTCTCTGGGAAATCGTGGA | qRTPCR |
| mmIL6R | GGTACTCCAGAAGACCAGAGGA | qRTPCR |
| mmIL10F | CCCAGAAATCAAGGAGC | qRTPCR |
| mmIL10R | TCACTCTTCACCTGCTCCAC | qRTPCR |
| mmIL1bF | CAGCAGCACATCAACAAG | qRTPCR |
| mmIL1bR | GTGCTCATGTCCTCATCCTG | qRTPCR |
| mmIL13F | GATCTGTGTCTCTCCCTCTGACCC | qRTPCR |
| mmIl13R | GCCTTGCGGTTACAGAGGCC | qRTPCR |
| mmMIP2AF | GAACAAAGGCAAGGCTAACTGA | qRTPCR |
| mmMIP2AR | AACATAACAACATCTGGGCAAT | qRTPCR |
| mmMxAF | GGCAGACACCACATACAACC | qRTPCR |
| mmMxAR | CCTCAGGCTAGATGGCAAG | qRTPCR |
| mmBeta Actin F | AATCGTGCGTGACATCAAAG | qRTPCR |
| mmBeta Actin R | ATGCCACAGGATTCCATACC | qRTPCR |
| Stat2F | GGATTCTGAATCAGGCTCAAAGAG | genotyping |
| Stat2R | GAGGTAAGAGGTTCCGAGTGTGTT | genotyping |
| Neo | CAGCGCATCGCCTTCTATCGCCTTCTTG | genotyping |
