## Supplemental Table 2 for "Stat2 loss disrupts damage signalling and is protective in acute pancreatitis"

**Supplemental Table 2. Mean quantitative normalised spectra for 348 unique phosphoproteins that were differently expressed in each experimental condition shown.**

| Protein symbol and name | Uniprot ID | WT PBS | WT + Cerulein | Stat2$^{-/-}$ | Stat2$^{-/-}$ + Cerulein |
|---|---|---|---|---|---|
| AGFG1 Arf-GAP domain and FG repeats-containing protein 1 | Q8K2K6 | 1.13 | 0.00 | 0.00 | 0.56 |
| LYRIC Protein LYRIC | Q80WJ7 | 1.33 | 0.00 | 0.22 | 1.23 |
| FKB15 FK506-binding protein 15 | Q6P9Q6 | 0.83 | 0.00 | 0.22 | 0.17 |
| NSUN2 tRNA (cytosine-5-)-methyltransferase NSUN2 | Q1HFZ0 | 1.38 | 0.00 | 0.64 | 0.73 |
| DHSO Sorbitol dehydrogenase | Q64442 | 1.38 | 0.00 | 0.64 | 0.63 |
| MRCKB Serine/threonine-protein kinase MRCK beta | Q7TT50 | 2.18 | 0.00 | 1.14 | 0.38 |
| EDC4 Enhancer of mRNA-decapping protein 4 | Q3UJB9 | 1.91 | 0.00 | 1.14 | 0.57 |
| KAP0 cAMP-dependent protein kinase type I-alpha regulatory subunit | Q9DBC7 | 1.55 | 0.00 | 1.04 | 1.01 |
| NDKB Nucleoside diphosphate kinase B | Q01768 | 0.83 | 0.00 | 1.11 | 0.60 |
| RGPA1 Ral GTPase-activating protein subunit alpha-1 | Q6GYP7 | 0.50 | 0.00 | 0.68 | 0.80 |
| TBCD4 TBC1 domain family member 4 | Q8BYJ6 | 0.51 | 0.00 | 0.71 | 0.17 |
| ITB4 Integrin beta-4 | A2A863 | 0.49 | 0.00 | 0.71 | 0.38 |
| DYN2 Dynamin-2 | P39054 | 0.83 | 0.00 | 1.21 | 0.60 |
| SSH3 Protein phosphatase Slingshot homolog 3 | Q8K330 | 0.50 | 0.00 | 0.93 | 0.62 |
| INADL InaD-like protein | Q63ZW7 | 0.82 | 0.00 | 1.82 | 0.20 |
| GLUC Glucagon | P55095 | 3.24 | 0.00 | 7.87 | 2.08 |
| DP13B DCC-interacting protein 13-beta | Q8K3G9 | 0.25 | 0.00 | 0.75 | 0.56 |
| RGNEF Rho-guanine nucleotide exchange factor | P97433 | 0.25 | 0.00 | 0.93 | 0.20 |
| LRBA Lipopolysaccharide-responsive and beige-like anchor protein | Q9ESE1 | 0.33 | 0.00 | 1.31 | 0.97 |
| RS6 40S ribosomal protein S6 | P62754 | 0.25 | 0.00 | 1.14 | 1.13 |
| CLAP2 CLIP-associating protein 2 | Q8BRT1 | 0.00 | 0.00 | 0.64 | 0.61 |
| RHG32 Rho GTPase-activating protein 32 | Q811P8 | 0.00 | 0.00 | 0.96 | 0.38 |
| RHG31 Rho GTPase-activating protein 31 | A6X8Z5 | 0.00 | 0.00 | 0.68 | 0.18 |

| | | | | | |
|---|---|---|---|---|---|
| SYTL5 Synaptotagmin-like protein 5 | Q80T23 | 0.00 | 0.00 | 1.14 | 0.24 |
| CTR2 Low affinity cationic amino acid transporter 2 | P18581 | 0.00 | 0.00 | 0.68 | 0.37 |
| G3P Glyceraldehyde-3-phosphate dehydrogenase | P16858 | 0.83 | 0.00 | 0.93 | 0.37 |
| SYNRG Synergin gamma | Q5SV85 | 0.83 | 0.00 | 0.93 | 0.80 |
| SPF45 Splicing factor 45 | Q8JZX4 | 0.83 | 0.00 | 0.93 | 0.80 |
| SC31A Protein transport protein Sec31A | Q3UPL0 | 0.83 | 0.00 | 0.78 | 0.75 |
| TOM70 Mitochondrial import receptor subunit TOM70 | Q9CZW5 | 0.83 | 0.00 | 0.93 | 0.37 |
| RS7 40S ribosomal protein S7 | P62082 | 0.83 | 0.00 | 0.93 | 0.36 |
| MINK1 Misshapen-like kinase 1 | Q9JM52 | 1.05 | 0.00 | 1.11 | 0.62 |
| RBP1 RalA-binding protein 1 | Q62172 | 1.08 | 0.00 | 0.96 | 0.36 |
| PURB Transcriptional activator protein Pur-beta | O35295 | 1.08 | 0.00 | 0.93 | 0.37 |
| SRC8 Src substrate cortactin | Q60598 | 1.33 | 0.00 | 1.68 | 0.80 |
| FAM21 WASH complex subunit FAM21 | Q6PGL7 | 1.91 | 0.00 | 2.43 | 0.97 |
| FAK1 Focal adhesion kinase 1 | P34152 | 0.00 | 0.00 | 0.00 | 0.80 |
| EF2 Elongation factor 2 | P58252 | 0.00 | 0.00 | 0.22 | 0.80 |
| PAG1 Phosphoprotein associated with glycosphingolipid-enriched microdomains 1 | Q3U1F9 | 0.50 | 0.00 | 0.39 | 0.80 |
| NOL8 Nucleolar protein 8 | Q3UHX0 | 0.00 | 0.00 | 0.00 | 0.80 |
| KS6A6 Ribosomal protein S6 kinase alpha-6 | Q7TPS0 | 0.00 | 0.00 | 0.28 | 0.80 |
| EPN2 Epsin-2 | Q8CHU3 | 0.91 | 0.00 | 0.75 | 0.80 |
| PP6R2 Serine/threonine-protein phosphatase 6 regulatory subunit 2 | Q8R3Q2 | 0.00 | 0.00 | 0.00 | 0.80 |
| NXP20 Protein Noxp20 | Q9D281 | 0.00 | 0.00 | 0.00 | 0.80 |
| ZC3HE Zinc finger CCCH domain-containing protein 14 | Q8BJ05 | 0.25 | 0.00 | 0.00 | 0.97 |
| HBB1 Hemoglobin subunit beta-1 | P02088 | 1.00 | 0.27 | 0.71 | 1.28 |
| CD2AP CD2-associated protein | Q9JLQ0 | 0.83 | 0.38 | 1.17 | 1.74 |
| SMAP Small acidic protein | Q9R0P4 | 0.84 | 0.34 | 1.82 | 1.43 |
| ES8L2 Epidermal growth factor receptor kinase substrate 8-like | Q99K30 | 0.33 | 0.27 | 0.00 | 1.01 |

| | | | | | |
|---|---|---|---|---|---|
| protein 2 | | | | | |
| THUM1THUMP domain-containing protein 1 | Q99J36 | 0.00 | 0.28 | 0.61 | 1.04 |
| ARHG7 Rho guanine nucleotide exchange factor 7 | Q9ES28 | 1.05 | 0.28 | 0.64 | 0.99 |
| EMAL3 Echinoderm microtubule-associated protein-like 3 | Q8VC03 | 0.83 | 0.28 | 1.28 | 0.99 |
| GTPB1 GTP-binding protein 1 | O08582 | 0.25 | 0.55 | 2.25 | 1.84 |
| MY18A Myosin-XVIIIa | Q9JMH9 | 0.00 | 0.34 | 0.93 | 1.10 |
| ARFG2 ADP-ribosylation factor GTPase-activating protein 2 | Q99K28 | 0.58 | 0.55 | 0.82 | 1.74 |
| PGRC1 Membrane-associated progesterone receptor component 1 | O55022 | 1.96 | 0.68 | 2.06 | 2.15 |
| MTUS1 Microtubule-associated tumor suppressor 1 homolog | Q5HZI1 | 1.63 | 0.28 | 1.39 | 0.85 |
| DESP Desmoplakin | E9Q557 | 2.29 | 1.08 | 2.54 | 3.22 |
| C2C2L C2 domain-containing protein 2-like | Q80X80 | 1.42 | 0.61 | 3.28 | 1.80 |
| STIM1 Stromal interaction molecule 1 | P70302 | 1.34 | 0.88 | 1.70 | 2.55 |
| PLPL7 Patatin-like phospholipase domain-containing protein 7 | A2AJ88 | 0.50 | 0.28 | 1.11 | 0.80 |
| GORS1 Golgi reassembly-stacking protein 1 | Q91X51 | 1.38 | 0.55 | 1.35 | 1.59 |
| ACE Angiotensin-converting enzyme | P09470 | 0.25 | 0.28 | 0.00 | 0.80 |
| M3K7 Mitogen-activated protein kinase kinase kinase 7 | Q62073 | 0.25 | 0.28 | 0.22 | 0.80 |
| SYMC Methionyl-tRNA synthetase, cytoplasmic | Q68FL6 | 0.75 | 0.34 | 0.93 | 0.97 |
| EIF3B Eukaryotic translation initiation factor 3 subunit B | Q8JZQ9 | 0.83 | 0.81 | 1.92 | 2.30 |
| CBS Cystathionine beta-synthase | Q91WT9 | 2.37 | 0.95 | 4.13 | 2.66 |
| TBC15 TBC1 domain family member 15 | Q9CXF4 | 1.46 | 0.65 | 0.99 | 1.72 |
| GOGA5 Golgin subfamily A member 5 | Q9QYE6 | 1.65 | 1.30 | 1.78 | 3.35 |
| PRP4B Serine/threonine-protein kinase PRP4 homolog | Q61136 | 0.75 | 0.38 | 1.42 | 0.97 |
| MTSS1 Metastasis suppressor protein 1 | Q8R1S4 | 0.00 | 0.65 | 0.18 | 1.57 |
| MIA3 Melanoma inhibitory activity protein 3 | Q8BI84 | 3.04 | 0.92 | 2.85 | 2.19 |
| ADA17 Disintegrin and metalloproteinase domain-containing protein 17 | Q9Z0F8 | 0.50 | 0.34 | 0.93 | 0.80 |
| RAIN Ras-interacting protein 1 | Q3U0S6 | 0.33 | 0.34 | 0.64 | 0.80 |

| | | | | | |
|---|---|---|---|---|---|
| MPRI Cation-independent mannose-6-phosphate receptor | Q07113 | 0.58 | 0.34 | 0.64 | 0.80 |
| PDIP3 Polymerase delta-interacting protein 3 | Q8BG81 | 0.25 | 0.34 | 0.39 | 0.80 |
| BYST Bystin | O54825 | 0.00 | 0.34 | 0.18 | 0.80 |
| NIBAN Protein Niban | Q3UW53 | 2.22 | 1.03 | 2.28 | 2.41 |
| FA83H Protein FAM83H | Q148V8 | 1.54 | 0.92 | 2.14 | 2.09 |
| IF2P Eukaryotic translation initiation factor 5B | Q05D44 | 6.18 | 2.85 | 5.70 | 6.39 |
| KLD7A Kelch domain-containing protein 7A | A2APT9 | 0.25 | 0.28 | 0.64 | 0.61 |
| CDK18 Cell division protein kinase 18 | Q04899 | 0.25 | 0.28 | 0.82 | 0.60 |
| IRF3 Interferon regulatory factor 3 | P70671 | 0.50 | 0.38 | 0.39 | 0.80 |
| MYPT1 Protein phosphatase 1 regulatory subunit 12A | Q9DBR7 | 1.84 | 2.04 | 2.75 | 4.10 |
| PWP1 Periodic tryptophan protein 1 homolog | Q99LL5 | 1.30 | 0.28 | 0.71 | 0.56 |
| SNP23 Synaptosomal-associated protein 23 | O09044 | 1.05 | 0.88 | 0.93 | 1.77 |
| TPD52 Tumor protein D52 | Q62393 | 0.25 | 1.64 | 0.43 | 3.13 |
| TIF1B Transcription intermediary factor 1-beta | Q62318 | 0.75 | 1.74 | 1.40 | 3.33 |
| SCG2 Secretogranin-2 | Q03517 | 1.08 | 0.61 | 0.79 | 1.17 |
| K1522 Uncharacterized protein KIAA1522 | A2A7S8 | 0.00 | 0.54 | 0.93 | 0.98 |
| CMGA Chromogranin-A | P26339 | 2.91 | 0.61 | 2.14 | 1.10 |
| MPRIP Myosin phosphatase Rho-interacting protein | P97434 | 0.25 | 0.55 | 0.82 | 0.98 |
| SRRT Serrate RNA effector molecule homolog | Q99MR6 | 1.63 | 0.65 | 0.68 | 1.16 |
| HS90B Heat shock protein HSP 90-beta | P11499 | 0.80 | 0.88 | 1.11 | 1.55 |
| KIF1B Kinesin-like protein KIF1B | Q60575 | 0.00 | 0.55 | 0.39 | 0.91 |
| CING Cingulin | P59242 | 1.08 | 0.34 | 0.68 | 0.56 |
| ARHGG Rho guanine nucleotide exchange factor 16 | Q3U5C8 | 0.58 | 0.34 | 0.68 | 0.56 |
| RBM25 RNA-binding protein 25 | B2RY56 | 1.87 | 0.61 | 1.11 | 0.97 |
| PININ Pinin | O35691 | 0.83 | 0.61 | 1.32 | 0.98 |
| PLEC Plectin | Q9QXS1 | 1.63 | 1.26 | 1.82 | 1.99 |

| | | | | | |
|---|---|---|---|---|---|
| F122AProtein FAM122A | Q9DB52 | 2.96 | 0.99 | 2.85 | 1.55 |
| UBP10 Ubiquitin carboxyl-terminal hydrolase 10 | P52479 | 2.71 | 0.61 | 3.24 | 0.94 |
| FA54B Protein FAM54B | Q9CWE0 | 2.68 | 0.88 | 1.81 | 1.33 |
| NCOR2 Nuclear receptor corepressor 2 | Q9WU42 | 0.84 | 0.38 | 0.71 | 0.56 |
| MEF2D Myocyte-specific enhancer factor 2D | Q63943 | 0.25 | 0.92 | 0.86 | 1.35 |
| F195B Protein FAM195B | Q3UGS4 | 0.00 | 0.55 | 0.00 | 0.80 |
| SDS3 Sin3 histone deacetylase corepressor complex component SDS3 | Q8BR65 | 0.00 | 0.55 | 0.00 | 0.80 |
| TOIP1 Torsin-1A-interacting protein 1 | Q921T2 | 0.00 | 0.55 | 0.00 | 0.80 |
| UBR4 E3 ubiquitin-protein ligase UBR4 | A2AN08 | 0.49 | 0.27 | 0.75 | 0.39 |
| SPTA2 Spectrin alpha chain, brain | P16546 | 0.00 | 0.65 | 1.54 | 0.94 |
| LARP4 La-related protein 4 | Q8BWW4 | 1.24 | 2.79 | 2.99 | 4.04 |
| RANB3 Ran-binding protein 3 | Q9CT10 | 0.88 | 0.38 | 0.18 | 0.54 |
| FND3A Fibronectin type-III domain-containing protein 3A | Q8BX90 | 4.34 | 2.52 | 2.49 | 3.64 |
| KI21A Kinesin-like protein KIF21A | Q9QXL2 | 0.33 | 0.99 | 0.46 | 1.43 |
| HNRPC Heterogeneous nuclear ribonucleoproteins C1/C2 | Q9Z204 | 0.33 | 0.98 | 0.28 | 1.42 |
| ASPC1 Tether containing UBX domain for GLUT4 | Q8VBT9 | 0.51 | 0.55 | 0.28 | 0.80 |
| SON Protein SON | Q9QX47 | 1.66 | 0.99 | 1.67 | 1.41 |
| NCK1 Cytoplasmic protein NCK1 | Q99M51 | 0.33 | 0.55 | 0.64 | 0.77 |
| RPB1 DNA-directed RNA polymerase II subunit RPB1 | P08775 | 0.55 | 0.27 | 0.00 | 0.38 |
| K1C18 Keratin, type I cytoskeletal 18 | P05784 | 4.34 | 4.26 | 5.55 | 5.96 |
| AP3D1 AP-3 complex subunit delta-1 | O54774 | 2.41 | 1.80 | 3.42 | 2.47 |
| PML Probable transcription factor PML | Q60953 | 2.43 | 0.71 | 1.11 | 0.98 |
| BI2L1 Brain-specific angiogenesis inhibitor 1-associated protein 2-like protein 1 | Q9DBJ3 | 1.08 | 2.79 | 2.53 | 3.79 |
| MAP2 Microtubule-associated protein 2 | P20357 | 3.09 | 1.37 | 4.39 | 1.85 |
| CALX Calnexin | P35564 | 1.93 | 2.48 | 2.99 | 3.36 |
| BA2L2 Protein BAT2-like 2 | Q3TLH4 | 0.83 | 0.28 | 0.93 | 0.37 |

| | | | | | |
|---|---|---|---|---|---|
| AKAP1 A-kinase anchor protein 1, mitochondrial | O08715 | 2.24 | 0.88 | 2.14 | 1.18 |
| ZDHC5 Probable palmitoyltransferase ZDHHC5 | Q8VDZ4 | 1.63 | 0.61 | 0.43 | 0.81 |
| EF1D Elongation factor 1-delta | P57776 | 6.58 | 4.46 | 8.76 | 5.86 |
| BIG3 Brefeldin A-inhibited guanine nucleotide-exchange protein 3 | Q3UGY8 | 1.65 | 2.74 | 2.49 | 3.54 |
| MKNK1 MAP kinase-interacting serine/threonine-protein kinase 1 | O08605 | 0.00 | 1.54 | 0.39 | 1.94 |
| AAK1AP2-associated protein kinase 1 | Q3UHJ0 | 1.34 | 1.87 | 2.03 | 2.35 |
| BAD Bcl2 antagonist of cell death | Q61337 | 1.33 | 0.88 | 2.21 | 1.11 |
| F195A Protein FAM195A | Q9CQB2 | 1.08 | 0.34 | 0.57 | 0.42 |
| SDPRSerum deprivation-response protein | Q63918 | 1.84 | 0.65 | 2.46 | 0.81 |
| CAMP3 Calmodulin-regulated spectrin-associated protein 3 | Q80VC9 | 0.00 | 2.42 | 0.89 | 2.92 |
| FARP2 FERM, RhoGEF and pleckstrin domain-containing protein 2 | Q91VS8 | 0.00 | 0.92 | 0.00 | 1.11 |
| IF4B Eukaryotic translation initiation factor 4B | Q8BGD9 | 1.16 | 1.81 | 1.85 | 2.18 |
| MYCB2 Probable E3 ubiquitin-protein ligase MYCBP2 | Q7TPH6 | 0.00 | 0.61 | 0.82 | 0.74 |
| AFAD Afadin | Q9QZQ1 | 1.38 | 3.72 | 3.35 | 4.47 |
| RHG01 Rho GTPase-activating protein 1 | Q5FWK3 | 0.00 | 0.98 | 0.00 | 1.18 |
| BIN1 Myc box-dependent-interacting protein 1 | O08539 | 3.79 | 2.79 | 3.49 | 3.25 |
| RCAS1 Receptor-binding cancer antigen expressed on SiSo cells | Q9D0V7 | 0.83 | 0.88 | 1.28 | 1.01 |
| DOCK7Dedicator of cytokinesis protein 7 | Q8R1A4 | 0.75 | 0.55 | 0.71 | 0.61 |
| JUN Transcription factor AP-1 | P05627 | 0.00 | 1.60 | 0.00 | 1.79 |
| SYTL1 Synaptotagmin-like protein 1 | Q99N80 | 0.25 | 1.26 | 0.18 | 1.41 |
| YAP1 Yorkie homolog | P46938 | 2.13 | 0.88 | 2.03 | 0.98 |
| LMNA Prelamin-A/C | P48678 | 0.00 | 1.44 | 0.64 | 1.59 |
| 4EBP1 Eukaryotic translation initiation factor 4E-binding protein 1 | Q60876 | 11.58 | 4.98 | 12.54 | 5.50 |
| PAXI Paxillin | Q8VI36 | 1.63 | 1.26 | 0.86 | 1.39 |
| NACAM Nascent polypeptide-associated complex subunit alpha, muscle-specific form | P70670 | 3.65 | 2.52 | 3.99 | 2.77 |
| LATS1 Serine/threonine-protein kinase LATS1 | Q8BYR2 | 1.39 | 0.55 | 1.04 | 0.60 |

| | | | | | |
|---|---|---|---|---|---|
| EF1B Elongation factor 1-beta | O70251 | 2.41 | 4.90 | 4.81 | 5.33 |
| IBTK Inhibitor of Bruton tyrosine kinase | Q6ZPR6 | 1.38 | 0.92 | 1.17 | 0.98 |
| GNL3 Guanine nucleotide-binding protein-like 3 | Q8CI11 | 2.46 | 0.88 | 1.00 | 0.93 |
| ODBA 2-oxoisovalerate dehydrogenase subunit alpha, mitochondrial | P50136 | 13.67 | 9.68 | 9.87 | 10.16 |
| COBL1 Cordon-bleu protein-like 1 | Q3UMF0 | 6.75 | 11.58 | 13.75 | 12.08 |
| TR150 Thyroid hormone receptor-associated protein 3 | Q569Z6 | 5.17 | 5.86 | 3.46 | 6.11 |
| DC1L1 Cytoplasmic dynein 1 light intermediate chain 1 | Q8R1Q8 | 0.58 | 2.17 | 0.39 | 2.24 |
| TISB Zinc finger protein 36, C3H1 type-like 1 | P23950 | 0.00 | 1.90 | 0.00 | 1.96 |
| HNRH1 Heterogeneous nuclear ribonucleoprotein H | O35737 | 0.83 | 1.91 | 1.11 | 1.96 |
| RIC1 Protein RIC1 homolog | Q69ZJ7 | 0.33 | 0.61 | 0.93 | 0.62 |
| ATX2L Ataxin-2-like protein | Q7TQH0 | 0.80 | 4.32 | 0.61 | 4.40 |
| RAF1 RAF proto-oncogene serine/threonine-protein kinase | Q99N57 | 0.83 | 1.77 | 1.39 | 1.80 |
| PAIRB Plasminogen activator inhibitor 1 RNA-binding protein | Q9CY58 | 0.83 | 5.00 | 1.14 | 5.05 |
| EHD2 EH domain-containing protein 2 | Q8BH64 | 0.50 | 0.98 | 1.14 | 0.99 |
| TCOF Treacle protein | O08784 | 0.80 | 1.16 | 0.68 | 1.17 |
| ABLM1 Actin-binding LIM protein 1 | Q8K4G5 | 2.21 | 1.81 | 1.32 | 1.81 |
| RBP2 E3 SUMO-protein ligase RanBP2 | Q9ERU9 | 0.00 | 0.98 | 0.18 | 0.98 |
| STIM2 Stromal interaction molecule 2 | P83093 | 1.29 | 0.54 | 1.57 | 0.53 |
| NUFP2 Nuclear fragile X mental retardation-interacting protein 2 | Q5F2E7 | 1.66 | 2.69 | 1.50 | 2.61 |
| SPTB2 Spectrin beta chain, brain 1 | Q62261 | 2.94 | 1.26 | 1.82 | 1.22 |
| REPS2 RalBP1-associated Eps domain-containing protein 2 | Q80XA6 | 1.38 | 0.82 | 0.68 | 0.80 |
| DBPA DNA-binding protein A | Q9JKB3 | 1.96 | 1.25 | 2.28 | 1.21 |
| PKN1 Serine/threonine-protein kinase N1 | P70268 | 1.38 | 0.65 | 0.50 | 0.62 |
| TB182 182 kDa tankyrase-1-binding protein | P58871 | 1.41 | 6.35 | 1.93 | 6.07 |
| MATR3 Matrin-3 | Q8K310 | 0.80 | 1.60 | 1.04 | 1.53 |
| AAKB15'-AMP-activated protein kinase subunit beta-1 | Q9R078 | 2.50 | 2.25 | 1.85 | 2.15 |

| | | | | | |
|---|---|---|---|---|---|
| PCBP1 Poly(rC)-binding protein 1 | P60335 | 1.01 | 6.71 | 2.06 | 6.21 |
| REPS1 RalBP1-associated Eps domain-containing protein 1 | O54916 | 1.91 | 0.82 | 1.32 | 0.74 |
| RRBP1 Ribosome-binding protein 1 | Q99PL5 | 0.00 | 0.88 | 0.00 | 0.80 |
| RS3A 40S ribosomal protein S3a | P97351 | 0.50 | 0.61 | 0.86 | 0.56 |
| ACINU Apoptotic chromatin condensation inducer in the nucleus | Q9JIX8 | 9.33 | 3.51 | 5.38 | 3.17 |
| MYH9 Myosin-9 | Q8VDD5 | 0.00 | 2.65 | 2.03 | 2.39 |
| ILF3 Interleukin enhancer-binding factor 3 | Q9Z1X4 | 1.68 | 0.88 | 0.64 | 0.80 |
| PCKGM Phosphoenolpyruvate carboxykinase [GTP], mitochondrial | Q8BH04 | 0.83 | 0.88 | 0.43 | 0.80 |
| ZFY19 Zinc finger FYVE domain-containing protein 19 | Q9DAZ9 | 0.25 | 0.88 | 0.68 | 0.80 |
| DC1I2 Cytoplasmic dynein 1 intermediate chain 2 | O88487 | 0.00 | 0.88 | 0.18 | 0.80 |
| RS17 40S ribosomal protein S17 | P63276 | 2.76 | 1.53 | 2.10 | 1.36 |
| EIF3G Eukaryotic translation initiation factor 3 subunit G | Q9Z1D1 | 4.05 | 2.42 | 3.10 | 2.15 |
| SEPT9 Septin-9 | Q80UG5 | 0.55 | 0.92 | 0.36 | 0.82 |
| NPMNucleophosmin | Q61937 | 7.61 | 3.78 | 4.57 | 3.30 |
| SMTL2 Smoothelin-like protein 2 | Q8CI12 | 0.50 | 0.28 | 0.93 | 0.24 |
| BIG2 Brefeldin A-inhibited guanine nucleotide-exchange protein 2 | A2A5R2 | 0.80 | 0.28 | 0.46 | 0.24 |
| TBA1B Tubulin alpha-1B chain | P05213 | 0.25 | 0.92 | 0.64 | 0.80 |
| SNW1 SNW domain-containing protein 1 | Q9CSN1 | 2.18 | 2.07 | 1.57 | 1.79 |
| IF4G3 Eukaryotic translation initiation factor 4 gamma 3 | Q80XI3 | 1.08 | 1.87 | 1.11 | 1.59 |
| GEPH Gephyrin | Q8BUV3 | 0.00 | 4.23 | 0.72 | 3.58 |
| CTGE5 Cutaneous T-cell lymphoma-associated antigen 5 homolog | Q8R311 | 6.07 | 6.47 | 3.99 | 5.46 |
| NUCKS Nuclear ubiquitous casein and cyclin-dependent kinases substrate | Q80XU3 | 2.79 | 4.39 | 3.49 | 3.66 |
| CHIP STIP1 homology and U box-containing protein 1 | Q9WUD1 | 2.43 | 1.91 | 1.18 | 1.59 |
| CGNL1 Cingulin-like protein 1 | Q6AW69 | 1.38 | 1.87 | 1.82 | 1.55 |
| CQ028 UPF0663 transmembrane protein C17orf28 homolog | Q8R1F6 | 2.46 | 3.27 | 2.14 | 2.70 |
| LIMA1 LIM domain and actin-binding protein 1 | Q9ERG0 | 2.17 | 0.92 | 1.04 | 0.76 |

| | | | | | |
|---|---|---|---|---|---|
| IF4G1 Eukaryotic translation initiation factor 4 gamma 1 | Q6NZJ6 | 1.63 | 4.44 | 1.03 | 3.66 |
| TOM1 Target of Myb protein 1 | O88746 | 0.00 | 1.80 | 0.18 | 1.47 |
| PACN3 Protein kinase C and casein kinase II substrate protein 3 | Q99JB8 | 0.25 | 0.98 | 0.53 | 0.80 |
| TTP Tristetraprolin | P22893 | 0.00 | 3.68 | 0.18 | 2.95 |
| CTNB1 Catenin beta-1 | Q02248 | 1.08 | 2.93 | 3.04 | 2.33 |
| VIGLN Vigilin | Q8VDJ3 | 1.96 | 2.70 | 1.35 | 2.15 |
| HDGF Hepatoma-derived growth factor | P51859 | 4.47 | 3.92 | 2.89 | 3.12 |
| RBM39 RNA-binding protein 39 | Q8VH51 | 3.80 | 2.14 | 2.46 | 1.70 |
| EIF3C Eukaryotic translation initiation factor 3 subunit C | Q8R1B4 | 2.79 | 1.49 | 2.07 | 1.18 |
| DJC12 DnaJ homolog subfamily C member 12 | Q9R022 | 0.00 | 1.26 | 0.82 | 0.99 |
| PAK2 Serine/threonine-protein kinase PAK 2 | Q8CIN4 | 2.39 | 3.44 | 2.89 | 2.69 |
| HSPB1 Heat shock protein beta-1 | P14602 | 1.55 | 2.52 | 1.96 | 1.96 |
| SYEP Bifunctional aminoacyl-tRNA synthetase | Q8CGC7 | 1.05 | 1.26 | 0.64 | 0.98 |
| UBXN1 UBX domain-containing protein 1 | Q922Y1 | 1.96 | 1.26 | 1.67 | 0.98 |
| EP15R Epidermal growth factor receptor substrate 15-like 1 | Q60902 | 0.58 | 1.26 | 0.28 | 0.98 |
| UBP14 Ubiquitin carboxyl-terminal hydrolase 14 | Q9JMA1 | 0.25 | 1.26 | 0.93 | 0.97 |
| RBM14 RNA-binding protein 14 | Q8C2Q3 | 2.68 | 1.26 | 2.07 | 0.97 |
| ODPA Pyruvate dehydrogenase E1 component subunit alpha, somatic form, mitochondrial | P35486 | 12.36 | 8.85 | 10.13 | 6.76 |
| SSFA2 Sperm-specific antigen 2 homolog | Q922B9 | 4.49 | 3.23 | 2.92 | 2.44 |
| I2BP2 Interferon regulatory factor 2-binding protein 2 | E9Q1P8 | 0.51 | 1.30 | 0.39 | 0.98 |
| MAVS Mitochondrial antiviral-signaling protein | Q8VCF0 | 0.00 | 1.81 | 0.00 | 1.35 |
| TOP2B DNA topoisomerase 2-beta | Q64511 | 1.88 | 2.18 | 1.67 | 1.61 |
| NSF1C NSFL1 cofactor p47 | Q9CZ44 | 8.89 | 5.48 | 6.53 | 4.05 |
| BORG5 Cdc42 effector protein 1 | Q91W92 | 2.49 | 2.58 | 3.63 | 1.91 |
| HSPB8 Heat shock protein beta-8 | Q9JK92 | 0.00 | 1.57 | 0.00 | 1.15 |
| SNIP1 Smad nuclear-interacting protein 1 | Q8BIZ6 | 2.17 | 2.42 | 1.71 | 1.77 |

| | | | | | |
|---|---|---|---|---|---|
| PKHA7 Pleckstrin homology domain-containing family A member 7 | Q3UIL6 | 0.25 | 0.27 | 1.11 | 0.20 |
| NCOR1 Nuclear receptor corepressor 1 | Q60974 | 0.58 | 0.27 | 0.64 | 0.20 |
| BCLF1 Bcl-2-associated transcription factor 1 | Q8K019 | 8.73 | 6.20 | 6.41 | 4.49 |
| RCAN1 Calcipressin-1 | Q9JHG6 | 0.00 | 4.43 | 0.00 | 3.20 |
| NOP58 Nucleolar protein 58 | Q6DFW4 | 4.39 | 2.42 | 2.31 | 1.73 |
| ARFG1ADP-ribosylation factor GTPase-activating protein 1 | Q9EPJ9 | 4.43 | 2.52 | 3.52 | 1.79 |
| SVIL Supervillin | Q8K4L3 | 1.63 | 2.25 | 1.28 | 1.59 |
| ABI1 Abl interactor 1 | Q8CBW3 | 2.21 | 2.52 | 2.56 | 1.75 |
| BCKD [3-methyl-2-oxobutanoate dehydrogenase [lipoamide]] kinase, mitochondrial | O55028 | 4.43 | 4.05 | 3.70 | 2.77 |
| SLTM SAFB-like transcription modulator | Q8CH25 | 0.25 | 0.27 | 1.49 | 0.18 |
| KLC4 Kinesin light chain 4 | Q9DBS5 | 3.54 | 3.50 | 2.96 | 2.39 |
| RHG17 Rho GTPase-activating protein 17 | Q3UIA2 | 1.13 | 2.18 | 0.53 | 1.48 |
| HS90A Heat shock protein HSP 90-alpha | P07901 | 0.58 | 1.16 | 0.64 | 0.79 |
| CHSP1 Calcium-regulated heat stable protein 1 | Q9CR86 | 0.00 | 2.89 | 0.18 | 1.96 |
| RPTOR Regulatory-associated protein of mTOR | Q8K4Q0 | 0.25 | 1.54 | 0.28 | 1.04 |
| PGM1Phosphoglucomutase-1 | Q9D0F9 | 1.38 | 1.54 | 1.17 | 1.04 |
| KLC3 Kinesin light chain 3 | Q91W40 | 1.93 | 0.92 | 0.68 | 0.62 |
| PGM2 Phosphoglucomutase-2 | Q7TSV4 | 6.66 | 6.26 | 4.17 | 4.20 |
| SL9A1 Sodium/hydrogen exchanger 1 | Q61165 | 0.25 | 1.53 | 0.46 | 1.01 |
| CTND1 Catenin delta-1 | P30999 | 1.96 | 2.62 | 2.49 | 1.73 |
| K2C8 Keratin, type II cytoskeletal 8 | P11679 | 2.68 | 4.35 | 3.16 | 2.88 |
| PDPK1 3-phosphoinositide-dependent protein kinase 1 | Q9Z2A0 | 0.50 | 0.55 | 0.93 | 0.36 |
| SRBS2 Sorbin and SH3 domain-containing protein 2 | Q3UTJ2 | 6.04 | 5.08 | 4.99 | 3.21 |
| FA40A Protein FAM40A | Q8C079 | 1.93 | 1.26 | 0.93 | 0.80 |
| FINC Fibronectin | P11276 | 1.63 | 1.26 | 0.93 | 0.80 |
| CCNY Cyclin-Y | Q8BGU5 | 2.76 | 2.52 | 1.85 | 1.59 |

| | | | | | |
|---|---|---|---|---|---|
| LRRF1 Leucine-rich repeat flightless-interacting protein 1 | Q3UZ39 | 0.91 | 1.26 | 0.61 | 0.80 |
| PI4KB Phosphatidylinositol 4-kinase beta | Q8BKC8 | 1.55 | 2.52 | 1.35 | 1.59 |
| DOCK1 Dedicator of cytokinesis protein 1 | Q8BUR4 | 0.25 | 1.26 | 0.18 | 0.80 |
| SYTL4 Synaptotagmin-like protein 4 | Q9R0Q1 | 0.00 | 1.26 | 0.46 | 0.80 |
| RBM26 RNA-binding protein 26 | Q6NZN0 | 1.33 | 2.14 | 2.07 | 1.35 |
| MAP4 Microtubule-associated protein 4 | P27546 | 0.00 | 2.14 | 0.00 | 1.35 |
| FLNB Filamin-B | Q80X90 | 0.00 | 2.51 | 0.00 | 1.58 |
| YBOX1 Nuclease-sensitive element-binding protein 1 | P62960 | 0.83 | 1.87 | 0.93 | 1.18 |
| LBH Protein LBH | Q9CX60 | 0.00 | 2.25 | 0.36 | 1.41 |
| BAT2 Large proline-rich protein BAT2 | Q7TSC1 | 0.83 | 1.20 | 0.93 | 0.74 |
| MAP7 Ensconsin | O88735 | 1.41 | 0.71 | 0.89 | 0.44 |
| S23IP SEC23-interacting protein | Q6NZC7 | 0.82 | 2.89 | 0.50 | 1.77 |
| RL24 60S ribosomal protein L24 | Q8BP67 | 0.00 | 1.20 | 0.00 | 0.73 |
| FETUA Alpha-2-HS-glycoprotein | P29699 | 22.88 | 34.42 | 20.03 | 20.43 |
| SC61B Protein transport protein Sec61 subunit beta | Q9CQS8 | 10.94 | 14.22 | 7.44 | 8.36 |
| TCEA1 Transcription elongation factor A protein 1 | P10711 | 4.64 | 5.17 | 4.28 | 2.95 |
| FCHO2 FCH domain only protein 2 | Q3UQN2 | 0.84 | 2.79 | 1.31 | 1.59 |
| MBB1A Myb-binding protein 1A | Q7TPV4 | 1.05 | 2.14 | 0.00 | 1.21 |
| RS20 40S ribosomal protein S20 | P60867 | 0.00 | 2.14 | 0.00 | 1.21 |
| PKP4 Plakophilin-4 | Q68FH0 | 0.58 | 2.51 | 1.67 | 1.42 |
| GSK3A Glycogen synthase kinase-3 alpha | Q2NL51 | 3.01 | 2.52 | 1.85 | 1.42 |
| ZO1 Tight junction protein ZO-1 | P39447 | 2.29 | 1.64 | 1.54 | 0.91 |
| GFPT1 Glucosamine--fructose-6-phosphate aminotransferase [isomerizing] 1 | P47856 | 4.14 | 2.89 | 1.60 | 1.59 |
| UBP8 Ubiquitin carboxyl-terminal hydrolase 8 | Q80U87 | 0.25 | 1.91 | 0.36 | 1.04 |
| RTN4 Reticulon-4 | Q99P72 | 1.34 | 1.81 | 0.18 | 0.98 |
| PERQ2 PERQ amino acid-rich with GYF domain-containing protein 2 | Q6Y7W8 | 1.30 | 1.81 | 1.07 | 0.97 |

| | | | | | |
|---|---|---|---|---|---|
| NDRG1 Protein NDRG1 | Q62433 | 2.13 | 1.81 | 1.72 | 0.97 |
| SEPT2 Septin-2 | P42208 | 1.66 | 1.26 | 2.03 | 0.68 |
| SARG Specifically androgen-regulated gene protein | Q8BI29 | 0.00 | 2.52 | 0.00 | 1.35 |
| CP013 UPF0585 protein C16orf13 homolog | Q9DCS2 | 1.41 | 1.87 | 1.39 | 0.99 |
| MUC1 Mucin-1 | Q02496 | 0.50 | 0.38 | 0.68 | 0.20 |
| EPIPL Epiplakin | Q8R0W0 | 0.83 | 1.53 | 0.93 | 0.80 |
| EVI1MDS1 and EVI1 complex locus protein EVI1 | P14404 | 0.50 | 0.34 | 0.28 | 0.17 |
| RS3 40S ribosomal protein S3 | P62908 | 2.13 | 3.85 | 1.60 | 1.96 |
| AASD1 Alanyl-tRNA editing protein Aarsd1 | Q3THG9 | 1.67 | 1.87 | 1.85 | 0.94 |
| LYST Lysosomal-trafficking regulator | P97412 | 0.50 | 1.26 | 0.93 | 0.62 |
| VINC Vinculin | Q64727 | 0.55 | 0.38 | 0.00 | 0.18 |
| ARHG1 Rho guanine nucleotide exchange factor 1 | Q61210 | 0.00 | 1.26 | 0.00 | 0.61 |
| SYNJ1 Synaptojanin-1 | Q8CHC4 | 0.00 | 0.88 | 0.00 | 0.42 |
| CCKAR Cholecystokinin receptor type A | O08786 | 0.00 | 1.15 | 0.22 | 0.55 |
| VASP Vasodilator-stimulated phosphoprotein | P70460 | 0.58 | 1.19 | 0.22 | 0.56 |
| ERRFI ERBB receptor feedback inhibitor 1 | Q99JZ7 | 0.00 | 1.26 | 0.00 | 0.57 |
| SCFD1 Sec1 family domain-containing protein 1 | Q8BRF7 | 2.51 | 1.77 | 1.60 | 0.80 |
| MTMR2 Myotubularin-related protein 2 | Q9Z2D1 | 0.50 | 0.98 | 1.32 | 0.42 |
| OSBL3 Oxysterol-binding protein-related protein 3 | Q9DBS9 | 0.25 | 0.98 | 0.46 | 0.42 |
| BAG3 BAG family molecular chaperone regulator 3 | Q9JLV1 | 2.21 | 0.99 | 1.11 | 0.42 |
| K1C19 Keratin, type I cytoskeletal 19 | P19001 | 0.51 | 2.41 | 0.89 | 0.98 |
| TPPP Tubulin polymerization-promoting protein | Q7TQD2 | 0.58 | 1.94 | 1.81 | 0.75 |
| ARK72 Aflatoxin B1 aldehyde reductase member 2 | Q8CG76 | 0.00 | 1.60 | 0.00 | 0.61 |
| ZCH18 Zinc finger CCCH domain-containing protein 18 | Q0P678 | 2.51 | 0.98 | 0.71 | 0.37 |
| IF4G2 Eukaryotic translation initiation factor 4 gamma 2 | Q62448 | 2.76 | 0.55 | 1.36 | 0.18 |
| CSRP1 Cysteine and glycine-rich protein 1 | P97315 | 0.83 | 1.26 | 0.18 | 0.42 |

| | | | | | |
|---|---|---|---|---|---|
| PP4R4 Serine/threonine-protein phosphatase 4 regulatory subunit 4 | Q8C0Y0 | 0.00 | 0.71 | 0.64 | 0.24 |
| COPB2 Coatomer subunit beta' | O55029 | 0.83 | 1.26 | 0.64 | 0.42 |
| WDR20 WD repeat-containing protein 20 | Q9D5R2 | 0.83 | 1.26 | 0.93 | 0.42 |
| GTF2I General transcription factor II-I | Q9ESZ8 | 0.25 | 0.61 | 0.68 | 0.17 |
| PDCD4 Programmed cell death protein 4 | Q61823 | 0.00 | 0.61 | 0.64 | 0.17 |
| NED4L E3 ubiquitin-protein ligase NEDD4-like | Q8CFI0 | 0.83 | 1.26 | 0.93 | 0.36 |
| NFX1 Transcriptional repressor NF-X1 | B1AY10 | 1.13 | 0.65 | 0.46 | 0.18 |
| ANR17 Ankyrin repeat domain-containing protein 17 | Q99NH0 | 0.00 | 0.88 | 0.00 | 0.24 |
| SYNP2 Synaptopodin-2 | Q91YE8 | 0.83 | 0.82 | 0.46 | 0.17 |
| ASPP2 Apoptosis-stimulating of p53 protein 2 | Q8CG79 | 0.00 | 1.26 | 0.00 | 0.24 |
| SNTB2 Beta-2-syntrophin | Q61235 | 0.55 | 1.26 | 0.00 | 0.20 |
| BSPRY B box and SPRY domain-containing protein | Q80YW5 | 0.00 | 1.36 | 0.18 | 0.20 |
| VP26B Vacuolar protein sorting-associated protein 26B | Q8C0E2 | 0.00 | 2.29 | 0.00 | 0.17 |
| STALP AMSH-like protease | Q76N33 | 0.83 | 0.00 | 0.00 | 0.00 |
| AT2A2 Sarcoplasmic/endoplasmic reticulum calcium ATPase 2 | O55143 | 1.93 | 0.38 | 0.25 | 0.00 |
| F125A Multivesicular body subunit 12A | Q78HU3 | 0.83 | 0.00 | 0.28 | 0.00 |
| PHC3 Polyhomeotic-like protein 3 | Q8CHP6 | 1.05 | 0.00 | 0.64 | 0.00 |
| MAST3 Microtubule-associated serine/threonine-protein kinase 3 | Q3U214 | 1.13 | 0.92 | 0.71 | 0.00 |
| ENSA Alpha-endosulfine | P60840 | 1.41 | 0.00 | 0.93 | 0.00 |
| KAP2 cAMP-dependent protein kinase type II-alpha regulatory subunit | P12367 | 1.38 | 0.88 | 0.93 | 0.00 |
| ANS1A Ankyrin repeat and SAM domain-containing protein 1A | P59672 | 1.32 | 0.27 | 0.93 | 0.00 |
| KLC2 Kinesin light chain 2 | O88448 | 0.58 | 0.27 | 0.93 | 0.00 |
| FA83F Protein FAM83F | Q3UKU4 | 0.25 | 0.00 | 0.68 | 0.00 |
| PH4H Phenylalanine-4-hydroxylase | P16331 | 0.25 | 0.00 | 0.86 | 0.00 |
| APCAdenomatous polyposis coli protein | Q61315 | 0.00 | 0.00 | 0.68 | 0.00 |
| FRY Protein furry homolog | E9Q8I9 | 0.00 | 0.00 | 0.71 | 0.00 |

| | | | | | |
|---|---|---|---|---|---|
| FYV1 1-phosphatidylinositol-3-phosphate 5-kinase | Q9Z1T6 | 0.00 | 0.00 | 0.93 | 0.00 |
| SCG3 Secretogranin-3 | P47867 | 0.00 | 0.34 | 0.93 | 0.00 |
| BRSK2BR serine/threonine-protein kinase 2 | Q69Z98 | 0.83 | 0.00 | 0.75 | 0.00 |
| UBP40 Ubiquitin carboxyl-terminal hydrolase 40 | Q8BWR4 | 0.83 | 0.00 | 0.71 | 0.00 |
| GALAGalanin | P47212 | 2.79 | 0.00 | 2.14 | 0.00 |
| U2AF2 Splicing factor U2AF 65 kDa subunit | P26369 | 1.38 | 0.34 | 1.11 | 0.00 |
| SCRIB Protein scribble homolog | Q80U72 | 1.08 | 0.27 | 1.00 | 0.00 |
| NDRG2 Protein NDRG2 | Q9QYG0 | 1.00 | 0.38 | 0.00 | 0.00 |
| NIPBL Nipped-B-like protein | Q6KCD5 | 0.25 | 0.38 | 0.43 | 0.00 |
| PAR3L Partitioning defective 3 homolog B | Q9CSB4 | 0.49 | 0.88 | 0.46 | 0.00 |
| K1C23 Keratin, type I cytoskeletal 23 | Q99PS0 | 0.25 | 0.61 | 0.22 | 0.00 |
| SASH1 SAM and SH3 domain-containing protein 1 | P59808 | 0.00 | 0.88 | 0.00 | 0.00 |
| HDAC4 Histone deacetylase 4 | Q6NZM9 | 0.00 | 0.98 | 0.00 | 0.00 |
| MP2K2Dual specificity mitogen-activated protein kinase kinase 2 | Q63932 | 0.00 | 0.99 | 0.00 | 0.00 |
| STALP AMSH-like protease | Q76N33 | 0.83 | 0.00 | 0.00 | 0.00 |
| AT2A2 Sarcoplasmic/endoplasmic reticulum calcium ATPase 2 | O55143 | 1.93 | 0.38 | 0.25 | 0.00 |
| F125A Multivesicular body subunit 12A | Q78HU3 | 0.83 | 0.00 | 0.28 | 0.00 |
| PHC3 Polyhomeotic-like protein 3 | Q8CHP6 | 1.05 | 0.00 | 0.64 | 0.00 |
| MAST3 Microtubule-associated serine/threonine-protein kinase 3 | Q3U214 | 1.13 | 0.92 | 0.71 | 0.00 |
| ENSA Alpha-endosulfine | P60840 | 1.41 | 0.00 | 0.93 | 0.00 |
| KAP2 cAMP-dependent protein kinase type II-alpha regulatory subunit | P12367 | 1.38 | 0.88 | 0.93 | 0.00 |
| ANS1A Ankyrin repeat and SAM domain-containing protein 1A | P59672 | 1.32 | 0.27 | 0.93 | 0.00 |
| KLC2 Kinesin light chain 2 | O88448 | 0.58 | 0.27 | 0.93 | 0.00 |
| FA83F Protein FAM83F | Q3UKU4 | 0.25 | 0.00 | 0.68 | 0.00 |
| PH4H Phenylalanine-4-hydroxylase | P16331 | 0.25 | 0.00 | 0.86 | 0.00 |
| APC Adenomatous polyposis coli protein | Q61315 | 0.00 | 0.00 | 0.68 | 0.00 |

| | | | | | |
|---|---|---|---|---|---|
| FRY Protein furry homolog | E9Q8I9 | 0.00 | 0.00 | 0.71 | 0.00 |
| FYV1 phosphatidylinositol-3-phosphate 5-kinase | Q9Z1T6 | 0.00 | 0.00 | 0.93 | 0.00 |
| SCG3 Secretogranin-3 | P47867 | 0.00 | 0.34 | 0.93 | 0.00 |
| BRSK2 BR serine/threonine-protein kinase 2 | Q69Z98 | 0.83 | 0.00 | 0.75 | 0.00 |
| UBP40 Ubiquitin carboxyl-terminal hydrolase 40 | Q8BWR4 | 0.83 | 0.00 | 0.71 | 0.00 |
| GALA Galanin | P47212 | 2.79 | 0.00 | 2.14 | 0.00 |
| U2AF2 Splicing factor U2AF 65 kDa subunit | P26369 | 1.38 | 0.34 | 1.11 | 0.00 |
| SCRIB Protein scribble homolog | Q80U72 | 1.08 | 0.27 | 1.00 | 0.00 |
| NDRG2 Protein NDRG2 | Q9QYG0 | 1.00 | 0.38 | 0.00 | 0.00 |
| NIPBL Nipped-B-like protein | Q6KCD5 | 0.25 | 0.38 | 0.43 | 0.00 |
| PAR3L Partitioning defective 3 homolog B | Q9CSB4 | 0.49 | 0.88 | 0.46 | 0.00 |
| K1C23 Keratin, type I cytoskeletal 23 | Q99PS0 | 0.25 | 0.61 | 0.22 | 0.00 |
| SASH1 SAM and SH3 domain-containing protein 1 | P59808 | 0.00 | 0.88 | 0.00 | 0.00 |
| HDAC4 Histone deacetylase 4 | Q6NZM9 | 0.00 | 0.98 | 0.00 | 0.00 |
| MP2K2 Dual specificity mitogen-activated protein kinase kinase 2 | Q63932 | 0.00 | 0.99 | 0.00 | 0.00 |
