## Supplemental Data for "Stat2 loss disrupts damage signalling and is protective in acute pancreatitis"

### Supplemental Fig 1

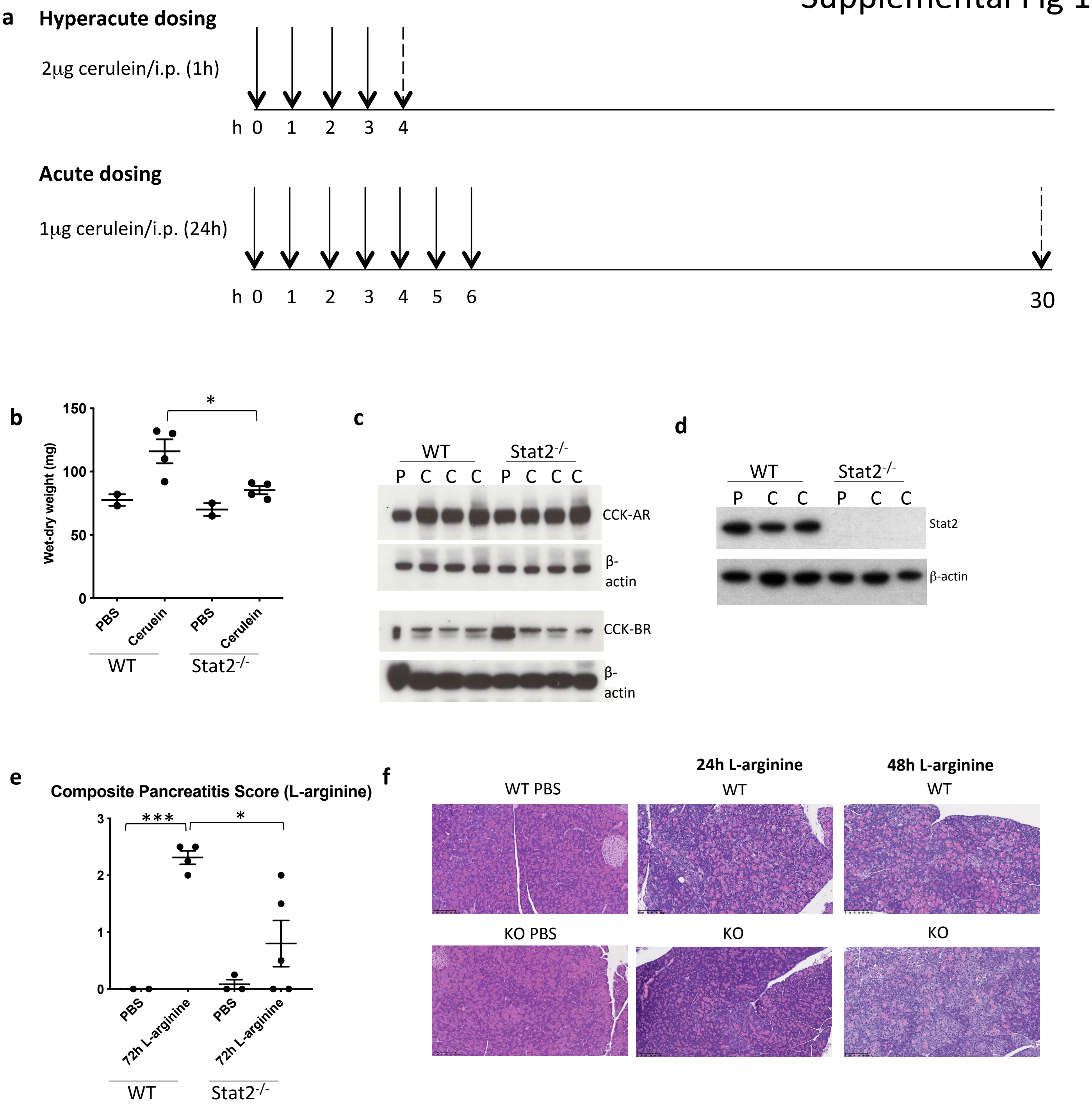

Supplemental Fig 2

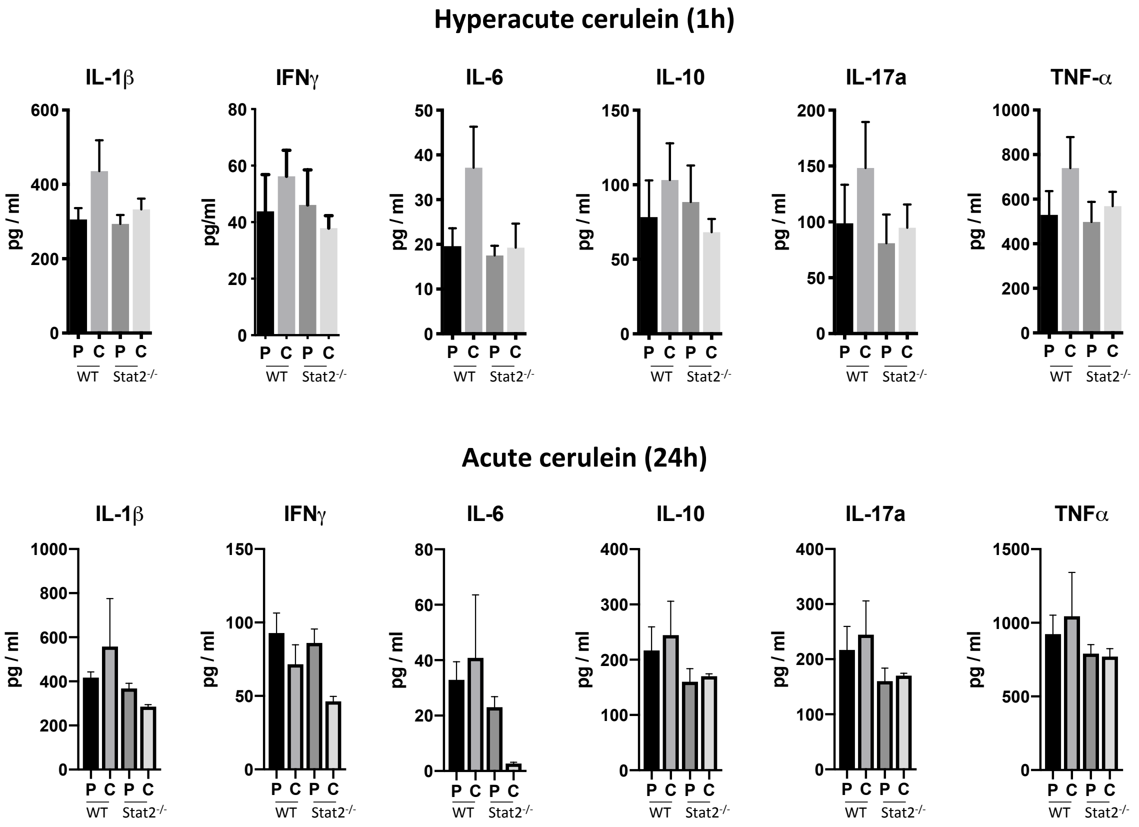

### Supplemental Figure 3

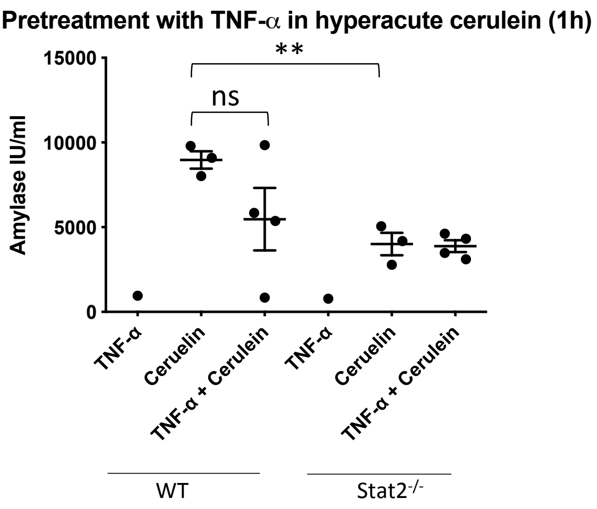

Supplemental Figure 4

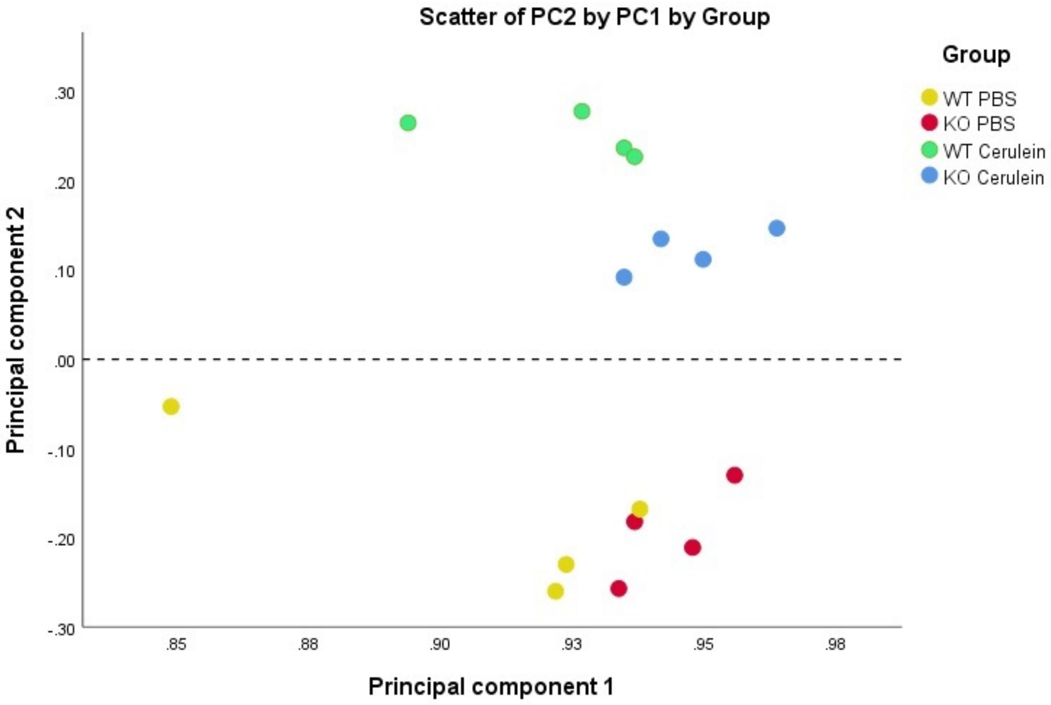

### Supplemental Figure 5

**a**

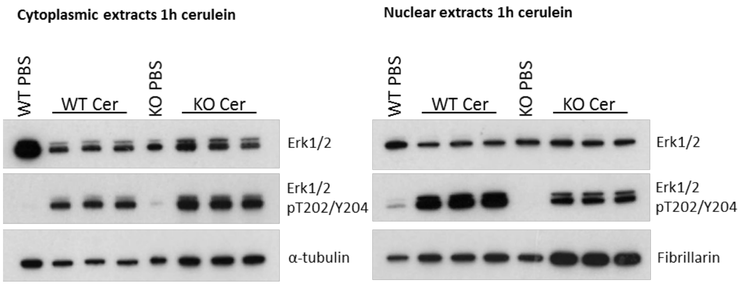


**b**

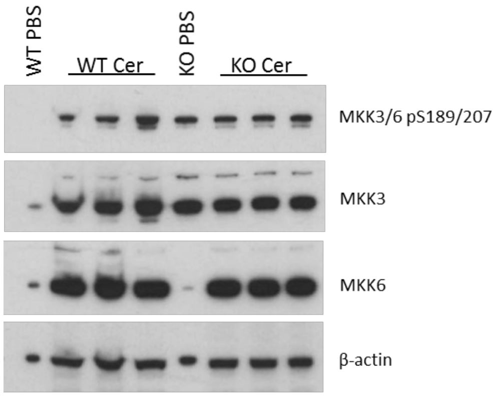
